# Theta-burst TMS of lateral occipital cortex reduces BOLD responses across category-selective areas in ventral temporal cortex

**DOI:** 10.1101/2020.08.03.230458

**Authors:** Iris I A Groen, Edward H Silson, David Pitcher, Chris I Baker

**Affiliations:** Section on Learning and Plasticity, Laboratory of Brain and Cognition, National Institute of Mental Health, Bethesda, MD 20892-1366, USA; Institute for Informatics, University of Amsterdam, Amsterdam, The Netherlands; Department of Psychology, School of Philosophy, Psychology and Language Sciences, The University of Edinburgh, Edinburgh, UK; Department of Psychology, The University of York, York, UK

**Author notes:** equal contribution (Iris I A Groen, Edward H Silson).

## Abstract

Human visual cortex contains three scene-selective regions in the lateral, medial and ventral cortex, termed the occipital place area (OPA), medial place area (MPA) and parahippocampal place area (PPA). Using functional magnetic resonance imaging (fMRI), all three regions respond more strongly when viewing visual scenes compared with isolated objects or faces. To determine how these regions are functionally and causally connected, we applied transcranial magnetic stimulation to OPA and measured fMRI responses before and after stimulation, using a theta-burst paradigm (TBS). To test for stimulus category-selectivity, we presented a range of visual categories (scenes, buildings, objects, faces). To test for specificity of any effects to TBS of OPA we employed two control conditions: Sham, with no TBS stimulation, and an active TBS-control with TBS to a proximal face-selective cortical region (occipital face area, or OFA). We predicted that TBS to OPA (but not OFA) would lead to decreased responses to scenes and buildings (but not other categories) in other scene-selective cortical regions. Across both ROI and whole-volume analyses, we observed decreased responses to scenes in PPA as a result of TBS. However, these effects were neither category specific, with decreased responses to all stimulus categories, nor limited to scene-selective regions, with decreases also observed in face-selective fusiform face area (FFA). Furthermore, similar effects were observed with TBS to OFA, thus effects were not specific to the stimulation site in the lateral occipital cortex. Whilst these data are suggestive of a causal, but non-specific relationship between lateral occipital and ventral temporal cortex, we discuss several factors that could have underpinned this result, such as the differences between TBS and online TMS, the role of anatomical distance between stimulated regions and how TMS effects are operationalised. Furthermore, our findings highlight the importance of active control conditions in brain stimulation experiments to accurately assess functional and causal connectivity between specific brain regions.

## Introduction

Category-selectivity is a fundamental organizing principle of high-level human visual cortex with regions selectively responsive to viewing faces (Kanwisher et al., 1997), scenes (Epstein & Kanwisher, 1998), objects (Malach et al., 1995) and bodies (Downing et al., 2001). For scenes, functional magnetic resonance imaging (fMRI) reveals a trio of selective regions, one on each of the lateral (Occipital Place Area, OPA), ventral (Parahippocampal Place Area, PPA) and medial (Medial Place Area, MPA) surfaces, respectively (Epstein & Kanwisher, 1998; Dilks et al., 2013; Silson et al., 2016). How these scene-selective regions are causally connected to form a putative scene-selective network and whether such interactions are specific to that network remains unclear. Here, we sought to test how visual information is routed through the scene-selective regions by combining transcranial magnetic stimulation (TMS) with consecutive fMRI, using a theta burst stimulation (TBS) paradigm.

Prior work employing TMS of category-selective regions has demonstrated the causal role they play in behavioral discriminations of stimuli from their preferred category (Pitcher et al., 2009; Dilks et al., 2013). These effects have been demonstrated in face-, body-, object- and more recently scene-selective regions in lateral occipital cortex (Pitcher et al., 2007; Dilks et al., 2013; Ganaden et al., 2013). These studies highlight the spatial- and stimulus-specificity that TMS can achieve when measured through behavioural performance on discrimination tasks. Beyond these local effects of stimulation on behavioral judgements of preferred categories, how stimulation of one category-selective region impacts information processing in other regions with the same category preference has only just started to be investigated.

In particular, recent work has explored this question with respect to the face-selective network using consecutive TMS/fMRI (Pitcher et al., 2014, 2017; Handwerker et al., 2020). Here, it was shown that TMS of the occipital face area (OFA; Gauthier et al., 2000) led to a local reduction in the response to faces within the OFA itself. Further, OFA stimulation also led to decreased responses in downstream face-selective regions, such as the fusiform face area (FFA) and the posterior superior temporal sulcus (pSTS - for dynamic faces). Similar findings of a local reduction in response, with some downstream effects have also been reported following TMS of scene-selective and object-selective regions (Mullin & Steeves, 2011; 2013; Rafique et al., 2015).

Here, we sought to investigate how causal intervention affects visual information processing across scene-selective regions. We compared TMS stimulation of OPA with both a no-stimulation control condition (Sham) and an active control condition (nearby face selective OFA) and measured BOLD responses to a range of stimulus categories across lateral occipital and ventral temporal cortex. We hypothesized that TMS of OPA, but not TMS of OFA or Sham, would lead to reduced response to scenes, but not other stimulus categories, in OPA as well as downstream scene-selective regions PPA and MPA.

## Methods and Materials

### 1. Participants and testing

A total of 16 participants completed the experiments (n=16, 12 females, mean age = 24.4 years). All participants had normal or corrected to normal vision and gave written informed consent. We tested 16 participants based on a survey of related studies. As far as we are aware no prior study had stimulated OPA with theta-burst stimulation making it hard to conduct a formal power analysis. Prior studies stimulating OPA and reporting effects measured with fMRI used a 1 Hz TMS protocol with sample sizes of 8 (Rafique et al., 2017) and 9 (Mullin et al., 2013) participants. By testing roughly twice as many participants and employing a rigorous within-participant design, we anticipated we would have sufficient power to detect fMRI effects of stimulation. This sample size is comparable to prior work focusing on face-selective OFA (Pitcher et al., 2014, n=15). The National Institutes of Health Institutional Review Board approved the consent and protocol. This work was supported by the Intramural Research program of the National Institutes of Health – National Institute of Mental Health Clinical Study Protocols 93-M-0170 and 12-M-0128, NCT00001360.

### 2. Data acquisition

Each participant completed four fMRI sessions: an initial functional localizer and population receptive field (pRF) mapping session, followed by three TMS/fMRI sessions: TMS to OPA, TMS to OFA (active control condition) and Sham (no-stimulation control condition). The localizer session was conducted first. Next, the order of the TMS/fMRI sessions was counterbalanced in a nested way, such that half of the participants received TMS to OPA before TMS to OFA, and half of the participants received Sham before active TMS (the full counterbalancing scheme is provided in **Supplementary Methods 1**.**0**). Fourteen participants completed the initial functional localisation session on the same 3.0T scanner as used for the TMS-fMRI experiments. The two remaining participants completed the functional localizer session on a 7.0T scanner as part of a different study (Groen et al., 2018).

#### 2.1 Scanning parameters 3.0T scanner

Initial functional localiser scans (n=14) and all TMS/fMRI scans (n=16) were conducted on a 3.0T GE Sigma MRI scanner in the Clinical Research Center on the National Institutes of Health campus (Bethesda, MD). Whole-brain volumes were acquired using an eight-channel head coil (28 slices; 3×3×4mm; 10% interslice gap; TR, 2 s, TE, 30ms; matrix size, 64×64, FOV, 192mm). In each TMS/fMRI scanning session, two T1-weighted anatomical images were acquired using the magnetization-prepared rapid gradient echo (MPRAGE) sequence (176 slices; 1×1×1mm; TR, 2.53 s, TE, 3.47 ms, TI, 900 ms, flip angle 7°).

#### 2.2 Scanning parameters 7.0T scanner

Initial functional localiser scans (n=2) were conducted on a 7.0T Siemens scanner (gradient echo planar imaging (EPI) sequence with a 32-channel head coil (47 slices; 1.6×1.6×1.6mm; 10% interslice gap; TR, 2 s; TE, 27 ms; matrix size, 126 × 126; FOV, 192 mm).

#### 2.3 Functional localizer paradigm

In fourteen participants, this session consisted of category localizer scans (6 runs). During category-localizer runs, color images from six categories (Scenes, Faces, Bodies, Buildings, Objects and Scrambled Objects) were presented at fixation in 16 s blocks (20 images per block, 300 ms per image, 500 ms blank), whilst participants performed a one-back task. Blocks were separated by 4 s blanks and started and ended with a 16 s baseline period. The total run length was 279 s (∼4.7 minutes). Each category was presented twice per run, with the order of presentation counterbalanced across participants and runs. Participants performed a one-back task. For the remaining two participants, initial localisation scans (4 runs) were completed on a 7.0T scanner. During category-localizer runs, grayscale images from eight categories (Scenes [man-made/natural, open/closed], Objects [man-made/natural], Faces and Buildings) were presented at fixation in 16 s blocks (20 images per block, 300 ms per image, 500 ms blank), whilst participants performed a one-back task (see Groen et al., 2018 for more details).

#### 2.4 pRF mapping paradigm

During pRF mapping, participants fixated centrally whilst a bar aperture traversed gradually throughout the visual field revealing scene fragments. During each run the bar aperture made eight sweeps through the visual field (4 directions, 2 orientations) and participants indicated a change in fixation color via a button press (see Silson et al., 2015 for more stimulus details). Fourteen participants completed six pRF mapping runs in the same session as the category localizer runs. For the two remaining participants, pRF mapping was conducted as part of a prior study employing the same paradigm (see Silson et al., 2015).

#### 2.5 Experimental TMS/fMRI sessions

Across the three TMS/fMRI sessions (OPA, OFA or Sham) participants fixated a central cross whilst grayscale images (10×10° visual angle) from eight different categories (Scenes [man-made/natural, open/closed], Objects [man-made/natural], Faces and Buildings) were back projected on a screen mounted onto the head coil with 1024×768 pixel resolution (**Figure 1A**). Images were presented in 16 s blocks (**Figure 1B)** (20 images per block, 300 ms per image, 500 ms grayscale background (blank) in between images). Consecutive blocks were separated by 8 s blank periods; in addition, each run started with and ended with a 16 s blank baseline period and included a 16 s baseline period in the middle of the run, resulting in a total run length of 415 s (∼6.9 minutes). Within each scanning session, participants completed 8 runs in total, 4 pre TMS and 4 Post TMS (see below), with T1-weighted anatomical scans occurring at the beginning and end of each session. The presentation order of categories were counterbalanced across each set of 4 runs such that each category was equally often (once) followed by every other category. To ensure that participants actively attended to the stimuli, participants were instructed to perform a 2-back task on the images by pressing a button on a hand-held button box, with stimulus repetitions occurring either 1, 2 or 3 times per block. Accuracy and reaction times were recorded.

**Figure 1:**
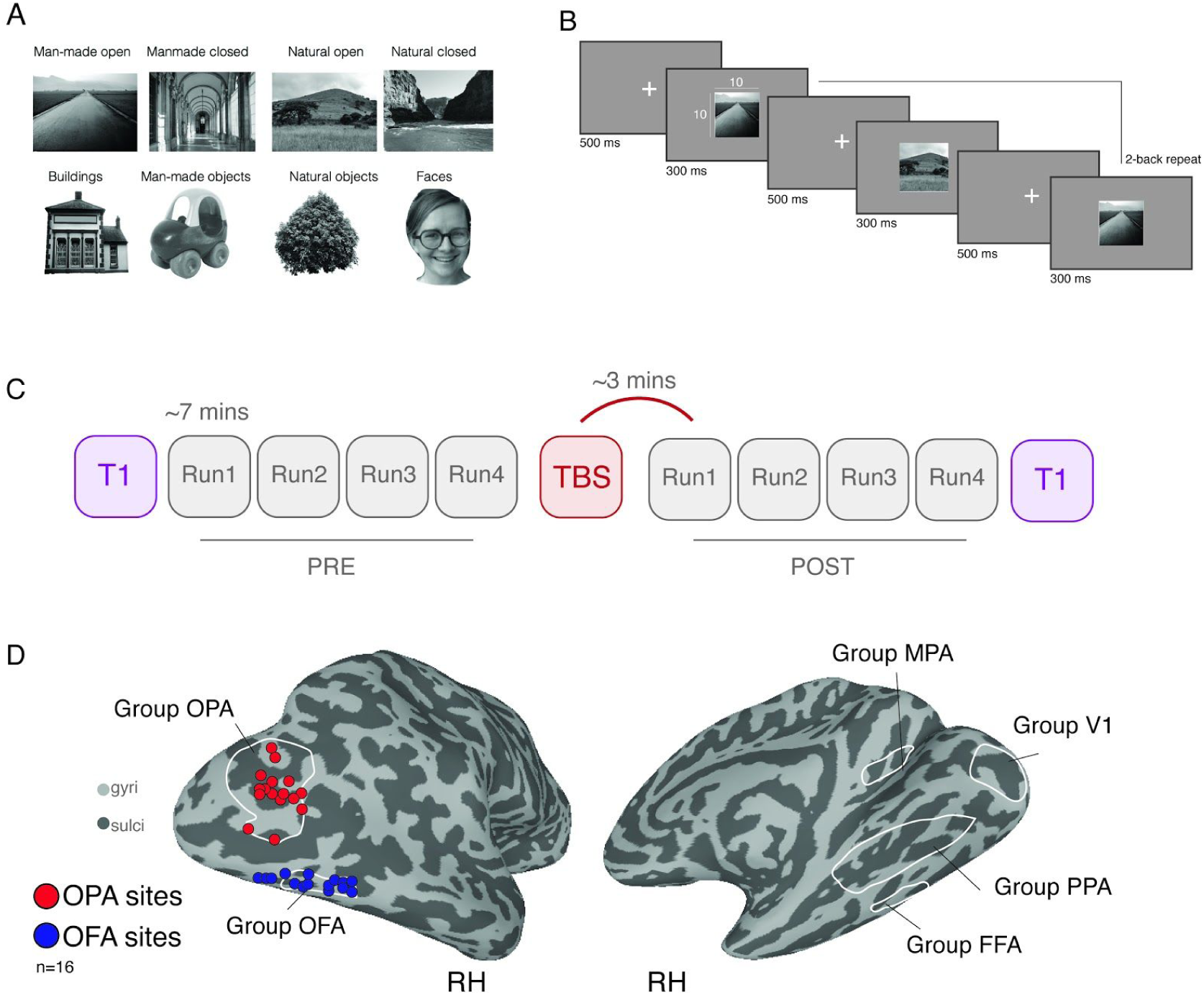
Example stimuli, task schematic, experimental design and TBS target sites. **A**, Example stimuli from eight categories: Four scene-categories (green outline) and four non-scene categories (blue outline). Note that the face exemplar has been omitted due to the copyright policy of the preprint server. **B**, Task schematic. Stimuli were presented in blocks. Participants fixated centrally and were required to push a button every time a 2-back repeat occurred. **C**, Schematic of experimental design. Each session began with a T1-weighted scan, followed by four Pre-TMS runs of the task, each lasting ∼7 mins. Following the end of Pre Run4, participants were removed from the scanner and TMS was performed. Four Post-TMS runs were then acquired followed by a second T1-weighted image. The first Post-TMS volumes were acquired ∼3 mins following the cessation of TBS. **D**, Left: Individual TMS stimulation sites (red = OPA, blue = OFA). Individual participant (n=16) stimulation sites are overlaid onto a partially inflated lateral view of the right hemisphere (gyri = light gray, sulci = dark gray). Right: Group ROIs included overlaid onto a medial view of the right hemisphere.

#### 2.6 TMS/fMRI experimental design

All three TMS/fMRI scanning sessions followed an identical design (**Figure 1C**). After initial head localization and slice prescription, a T-1 weighted anatomical image was acquired, followed by four Pre-TMS task runs. Immediately following the fourth Pre-TMS run, participants were removed from the scanner and seated on a stool inside the MRI control room. Once seated, the participants’ head was co-registered to their high-resolution structural scan using the BRAINSIGHT neuronavigation software (ROGUE research). During active TMS sessions, participants received 60 s of theta-burst stimulation to either the right OPA or right OFA. The position of the participants’ head and the stimulating coil were tracked in real-time and displacement errors were kept < 1mm. For the Sham session, the same procedure of first coregistration and then coil placement was performed, but no TMS pulses were delivered. A secondary coil, positioned close to, but away from the head, was used to produce the stereotypical noises associated with TMS. Immediately following cessation of TMS (or Sham), participants were returned to the MRI scanner. Participants then completed four Post-TMS task runs, followed by an additional T-1 weighted anatomical image. Post-TMS scanning commenced for all participants within 4 mins following TMS.

#### 2.7 TMS stimulation parameters

A Magstim Super Rapid 2 stimulator (Magstim, Wales, UK) was used to deliver TBS using a figure-of-eight coil (70 mm wing diameter). TBS was delivered at 30% of maximum machine output over the target voxel for each session. A continuous TBS paradigm (Huang et al., 2005) was used consisting of 3 pulses at 50 Hz repeated every 200 ms for a total of 900 pulses (60 s). This protocol was identical to that used in previous TMS/fMRI studies at NIH (Pitcher et al., 2014, 2017; Handwerker et al., 2020).

### 3. fMRI data pre-processing

All anatomical and functional data were pre-processed and analyzed using AFNI (Cox, 1996) (RRID: <u>SCR_005927</u>). Below we outline the preprocessing steps taken for both the initial functional localization and for the TMS/fMRI experiments.

#### 3.1 Functional localizer session

All images were motion corrected to the first volume of the first run (using the AFNI function *3dVolreg*) after removal of the appropriate dummy volumes to allow stabilization of the magnetic field. Following motion correction, images were spatially smoothed (*3dmerge*) with a 5 mm full-width-half-maximum smoothing kernel. Signal amplitudes were then converted into percent signal change (*3dTstat*). We employed a general linear model implemented in AFNI (*3dDeconvolve, 3dREMLfit*). The data at each time point were treated as the sum of all effects thought to be present at that time and the time-series was compared against a Generalized Least Squares (GSLQ) model fit with REML estimation of the temporal auto-correlation structure. Responses were modelled by convolving a standard gamma function with a 16 s square wave for each stimulus block. Estimated motion parameters were included as additional regressors of no-interest and fourth-order polynomials were included to account for slow drifts in the MR signal over time. To derive response magnitude per category, *t*-tests were performed between the category-specific beta coefficients and baseline.

#### 3.2 Population receptive field session

All images were motion corrected (*3dVolreg*) and detrended (*3dDetrend*) before being averaged across runs. The average time-series data were analysed using a 2-dimensional Gaussian pRF model implemented in AFNI. A detailed description of the model is available elsewhere (Silson et al., 2015). Briefly, given the position of the stimulus in the visual field at every time point, the model estimates the pRF parameters that yield the best fit to the data: pRF center location (x, y), and size (diameter of the pRF). Both Simplex and Powell optimization algorithms are used simultaneously to find the best time-series/parameter sets (x, y, size) by minimizing the least-squares error of the predicted time-series with the acquired time-series for each voxel.

#### 3.2 Functional TMS/fMRI sessions

Functional data from TMS sessions followed a similar pre-processing pipeline to that specified above, but differed in the following ways. For each TBS/fMRI session a mean anatomical image was first computed across the two T1 scans acquired before (Pre) and after (Post) TMS (*3dcalc*). This anatomical image is assumed to be half-way between the anatomical images acquired at the start and end of each session and therefore constitutes a reference image that is unbiased to either the Pre or the Post session. Once pre-processed, all EPI data within a session were then deobliqued (*3dWarp*) and aligned to this mean anatomical image (*align_epi_anat*.*py*). GLMs were estimated for each run separately (*3dDeconvolve, 3dREMLfit*) as opposed to concatenating all runs together before statistical analysis (default option in AFNI) in the unaligned, native volume space, after which the resulting statistical parametric maps were aligned to the mean anatomical image by applying the transformation matrices from the EPI alignment.

#### 3.3 TMS stimulation sites and ROI localization

Scene- and face-selective ROIs (OPA, OFA, PPA, FFA, MPA, **Figure 1D**) derived from the independent localizer session, were defined by the contrast of Scenes *versus* Faces (*p* < 0.0001, uncorrected). Stimulation sites for subsequent TMS sessions were identified on an individual participant basis using the Brainsight TMS-MRI co-registration system. In each participant, the statistical maps reflecting the contrast of Scenes *versus* Faces were overlaid onto the high-resolution anatomical scan. The primary OPA stimulation site was defined as the peak voxel of scene-selectivity within right OPA, and the active-control OFA stimulation site as the peak voxel of face-selectivity within right OFA. Individual target sites for each participant are depicted in **Figure 1D**. In addition, a control ROI in V1 was defined using the group-average pRF data encompassing the visual field representation subtended by the stimuli (**Figure 1D**).

#### 3.4 TMS accuracy

We took advantage of two measurements (coil-target error, coil-target distance; (see **Supplementary Figure S1, A-B**) automatically acquired through the Brainsight system that provide indices of stimulation precision (Silson et al., 2013). *Coil-target error* provides the minimum Euclidean distance between the projected vector of TMS and the sphere centred on the target voxel. We calculated the mean coil-target error in each TMS condition and submitted these to a linear-mixed model to test for a main effect of TMS condition (OPA, OFA & Sham). Neither the effect of TMS condition (F(2, 30)=1.98, p=0.15) nor any pairwise comparisons were significant (p>0.05). The lack of a TMS condition effect indicates an equivalent level of precision across the three conditions. This is valuable data as the Sham condition targeted the OPA and thus no difference between these conditions (OPA, Sham) would be expected. *Coil-target distance* measures the Euclidean distance between the target voxel and the calibration point of the TMS coil. For each TMS condition (OPA, OFA & Sham) we calculated the mean coil-target distance in each participant and submitted these to a linear-mixed model with TMS condition as the only main effect. Neither the main effect of TMS condition (F(2, 28.97)=1.02, p=0.37) nor any pairwise-comparisons (p>0.05) were significant. Although it is possible that a given target is further away (i.e. deeper in a sulcus) than another, the lack of a significant TMS condition effect suggests a) that our target sites were similarly distanced from the coil and b) that our calibration procedure was consistent across TMS sessions.

### 4. fMRI data analysis

#### 4.1 Sampling of data to the cortical surface

In each participant, the pre-processed functional data from all sessions were projected onto surface reconstructions (*3dvol2surf*) of each individual participant’s hemispheres derived from the Freesurfer4 autorecon script (http://surfer.nmr.mgh.harvard.edu/) using the Surface Mapping with AFNI (SUMA) software The freesurfer reconstructions were based on the T1s obtained in the localizer session. In order to align the functional data to these surfaces, the mean (Pre-Post) T1 from each TMS/fMRI session was first aligned to the volume used for surface reconstruction (*@SUMA_AlignToExperiment*).

#### 4.2 ROI definitions

ROIs were defined by overlaying the statistical results of the contrast Scenes versus Faces (taken from the initial functional localiser session) onto the surface reconstructions of each individual participant, before thresholding (p<0.0001, uncorrected). ROIs were defined using the interactive ROI drawing tool in SUMA according to both these statistical criteria and accepted anatomical locations (e.g. OPA is both scene-selective and overlaps with the transverse occipital sulcus). No further anatomical or functional constraints were applied.

#### 4.3 ROI analysis

Once defined, the vertices comprising these ROIs were converted to a 1D index of node indices per ROI (*ROI2dataset*), which was subsequently used to extract beta-coefficients for each stimulus category from the three separate TMS/fMRI sessions for each surface node within the ROI (*ConvertDset*). We verified that the localizer ROIs were adequately aligned with the experimental runs by visual inspection and by computing the Euclidean distance between the centers of mass of the localizer ROI and the Pre-TMS runs in each individual participant (see **Supplementary Figure S1, C-E)**. The extracted beta-coefficients were then imported into Matlab (Version R2018B) and averaged across all nodes within each ROI. The resulting ROI activation measures were fitted with a linear mixed effects model, using the *lme4* package implemented in R (Bates et al., 2015). The model comprised four fixed effects, one for each within-subject factor: TMS condition (OPA, OFA or Sham), Session (Pre, Post), Run (Runs 1-4) and Stimulus (Scenes [man-made/natural, open/closed], Objects [man-made/natural], Faces and Buildings). The estimated coefficients for each of the fixed effects were evaluated with omnibus ANOVAs, and the resulting p-values were corrected for multiple comparisons across the six examined ROIs (OPA, OFA, PPA, FFA, MPA, V1) using Bonferroni correction. The full statistical output for each ROI is provided in **Supplementary Results 1**, and the ROI data tables and R scripts used for the analyses are provided on the Open Science Framework (OSF) project page accompanying this paper (https://osf.io/6nq7r/). Our primary analyses modelled participants as a random effect using a random intercept term. Additional analyses included random effects for the within-subject factors (i.e. random slopes), and formal model comparison using Likelihood Ratio Tests (see **Supplementary Results 2**). Since TMS was applied to the right hemisphere, analyses were restricted to right hemisphere ROIs.

#### 4.4 Whole brain analysis

To investigate whole brain effects of TMS we employed a linear mixed effects model in AFNI (3dLME; Chen et al., 2013) in each hemisphere separately, although we focus primarily on the stimulated hemisphere (right). The model comprised the same four fixed effects as the ROI analyses (TMS condition, Session, Run and Stimulus). The model assessed the presence of significant main effects and their interactions. Our primary 3dLME analyses modelled participants as a random effect using a random intercept term. Based on the ROI model comparison analyses, additional whole-brain analyses also included random effects for the within-subject factors TMS condition and Session using a random slope (see **Supplementary Figure S5**). Whole brain results are provided on the OSF (https://osf.io/6nq7r/).

## Results

### Consecutive TMS/fMRI in scene-selective cortex: hypothesis and predictions

To assess the causal and functional relationships between scene-selective regions, fMRI BOLD responses to scene, building, face and object stimuli were measured in OPA, PPA and MPA before and after application of TMS to OPA. To control for non-TBS induced effects, the same participants also took part in an additional session in which the exact same procedure was followed but no TMS was applied (Sham; see Methods). Moreover, to determine the specificity of TMS to the scene network, we also included an active control session in which TMS was applied to a non scene-selective, but proximal region, the face-selective OFA.

As outlined above, we hypothesized that, relative to a baseline measurement taken prior to TMS stimulation, TMS to OPA would result in a reduced response in OPA itself as well as downstream scene-selective regions. If present, this reduced response could be category-specific in two ways: stimulus-specific, i.e. occurring for scene and building stimuli, but not faces or objects; and stimulation site-specific, i.e. occurring only as a result of stimulating scene-selective OPA, but not when stimulating face-selective OFA, which is thought to be part of a different category-selective network. These two types of category-specificity could be revealed by our experimental design in the following ways. First, if TMS effects are both stimulus- and site-specific, we should observe a decreased response to scene and building stimuli (but not face or object stimuli), in scene-selective regions as a result of TMS to OPA (but not OFA). Second, if TMS effects are site-specific but not stimulus-specific, stimulation to OPA (but not OFA) will cause scene-selective regions to respond less to *all* stimuli. A third possible scenario is that the TMS effects are neither stimulus-nor site-specific, such that stimulation to either OPA or OFA decreases responses across all stimuli.

We assessed the presence and category-specificity of TMS-induced response reductions in each scene-selective ROI using a linear mixed effects model (see Methods) that tested for fixed effects of TMS-condition (OPA, OFA, Sham), Session (Pre- or Post-TMS), Stimulus (Scenes [man-made/natural, open/closed], Objects [man-made/natural], Faces and Buildings). To take into account subject-specific difference in fMRI responses across ROIs, the model included subject as a random factor, and since prior reports have shown that TBS effects vary over time relative to stimulation (Pitcher et al., 2014; 2017), we additionally included Run (1-4) as a fixed effect. In this model, support for both site- and stimulus-specificity of TMS-induced effects would be evident through a significant interaction between TMS-condition, Session and Stimulus, such that responses selectively decrease as a result of TBS to OPA, for Scene stimuli, in the Post session. Support for site-specificity, but not stimulus-specificity would be evidenced by a two-way interaction between Session and TMS-condition (but not Stimulus), with effects of TBS for OPA but not OFA or Sham. Finally, a non-specific effect of TMS would be evident through a two-way interaction between Session and TMS-condition reflecting reductions as a result of stimulating both OPA and OFA but not Sham.

### ROI analysis

In OPA, contrary to our predictions, we found no evidence for an interaction between TMS-condition, Session and Stimulus (F(14, 2857) = 0.61, p = 0.86), nor any two-way interactions (full statistical reports for all ROIs can be found in **Supplementary Results 1**). A significant main effect was found for Stimulus (F(7, 2857) = 246.1, p < 0.001), reflecting the well-known category-selectivity of OPA; as shown in **Figure 2A**, the OPA shows the strongest response to scenes, followed by buildings and objects, and the weakest response for faces. These response patterns were strikingly consistent across the three TMS conditions and the Pre and Post sessions, indicating a lack of TMS effect on average across this ROI. This is further demonstrated in **Figure 2B**, which shows the response difference between the Pre and Post measurement for each condition: no significant reduction is observed.

**Figure 2:**
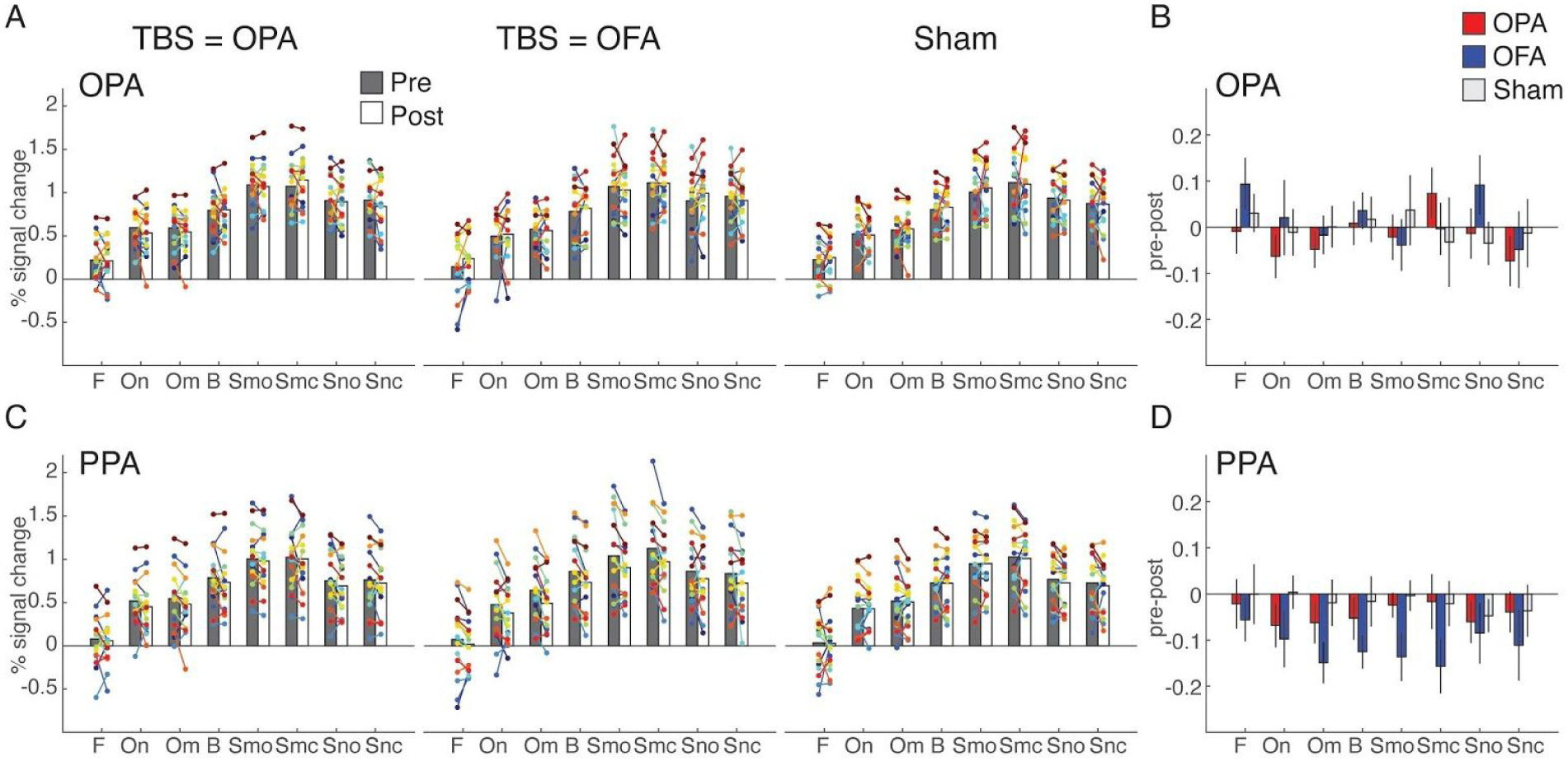
TMS effects in OPA and PPA. **A**, Bars represent the group-average response in OPA for each of the stimulus category blocks presented in the experiment (F = faces, On = natural objects, Om = man-made objects, B = buildings, Smo = Manmade open scenes, Smc = manmade closed scenes, Sno = natural open scenes, Snc = closed natural scenes) during Pre (gray) and Post (white) sessions measured during TMS to OPA (left), TMS to OFA (middle) and Sham (right). Individual participant data points are overlaid and linked in each case. **B**, Bars represent the response difference between the Pre and Post session for TMS to OPA (red), TMS to OFA (blue) and Sham (gray). Error bars represent S.E.M. across subjects. **C-D**, same as A-B, but for PPA.

In PPA, similarly no evidence was found for a three-way interaction between TMS-condition, Session and Stimulus (F(14, 210) = 0.26, p = 1; **Figure 2C**), but there was a significant two-way interaction of Session and TMS-condition (F(2,2857) = 9.1, p < 0.001, Bonferroni-corrected). As shown in **Figure 2D**, this interaction reflected a reduced response across stimulus categories as a result of stimulating OFA and OPA relative to Sham stimulation, although the effects were numerically stronger for stimulating OFA.

In MPA, no significant interactions related to TMS stimulation were observed, even though this region exhibited robust visual responses to scene stimuli in accordance with its category preference (see **Supplementary Figure S2**).

To investigate whether the observed pattern of results was specific to scene-selective regions, we additionally tested for TMS effects in face-selective OFA (i.e. the active control site) and the FFA. Interestingly, these ROIs showed a similar pattern as for the scene-selective regions: a lack of a local TMS effect at the stimulation ROI (OFA; **Figure 3A-B**) coupled with a significant two-way interaction of Session and TMS-condition (F(2,2857) = 11.3, p < 0.001, Bonferroni-corrected) in the downstream region (FFA; **Figure 3C-D**) indicating a reduced response across all stimuli as a result of TMS.

**Figure 3:**
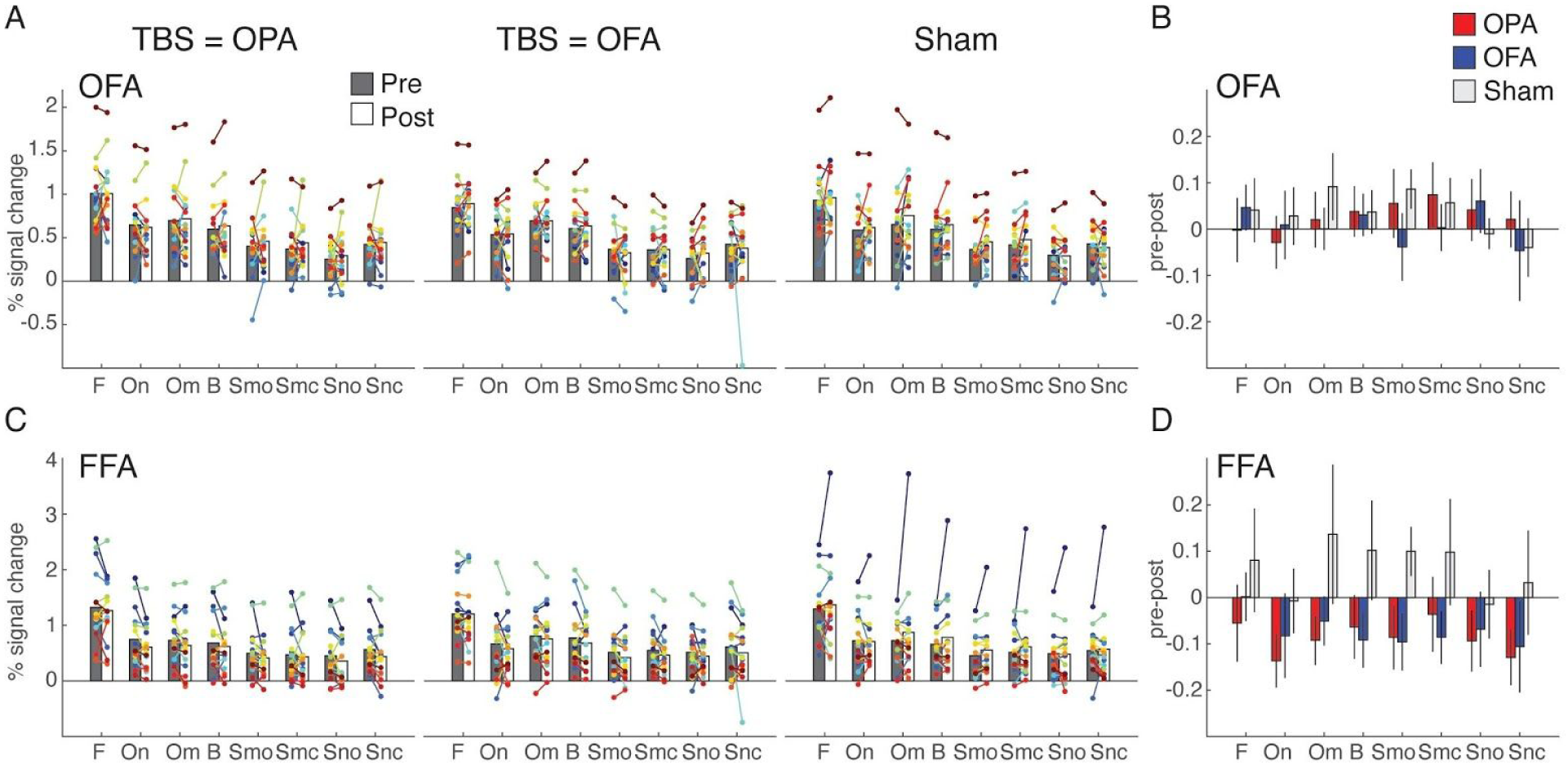
TMS effects in OFA and FFA. **A**, Bars represent the group-average response in OFA for each of the stimulus category blocks presented in the experiment (F = faces, On = natural objects, Om = man-made objects, B = buildings, Smo = Manmade open scenes, Smc = manmade closed scenes, Sno = natural open scenes, Snc = closed natural scenes) during Pre (gray) and Post (white) sessions measured during TMS to OPA (left), TMS to OFA (middle) and Sham (right). Individual participant data points are overlaid and linked in each case. **B**, Bars represent the response difference between the Pre and Post session for TMS to OPA (red), TMS to OFA (blue) and Sham (gray). Error bars represent S.E.M. across subjects. **C-D**, same as A-B, but for FFA. Note that for the Sham condition in FFA, the apparent increase in the Post relative to the Pre session is largely driven by one outlier subject. Statistical results were qualitatively the same when excluding this subject from analysis.

To examine potential effects of TMS outside category-selective cortex, we also examined responses in retinotopically defined V1 corresponding to the stimulus size (see Methods and **Supplementary Figure S2**). As expected, we observed robust visual responses to all stimulus categories, but no interaction effects related to TMS.

Finally, while none of the analyses indicated any interactions with Run indicative of a temporally specific TMS effect, a strong main effect of Run (all F(3,2857) > 41.1, all p < 0.0001) was found in all ROIs; further data inspection indicated that this main effect reflected an overall decrease in response over time within both Pre and Post sessions.

Taken together, the ROI analyses demonstrate effects of TMS to lateral occipital cortex in both scene- and face-selective downstream ventral temporal regions. However, these effects were neither stimulus-nor site-specific. While the reduced responses appear robust in ventral regions, especially for OFA stimulation, one puzzling aspect of the results is the lack of reduced activity in the stimulated lateral occipital ROIs. One possibility is that we overlooked a local effect within the stimulated ROIs themselves. To further clarify this pattern, we next proceeded to fit a whole-brain linear mixed effects model that is not constrained to average effects across large cortical regions.

### Whole brain analysis

Our initial analyses were based on the average responses within individually specified ROIs. Although principled, this approach involves averaging across large numbers of surface nodes in each ROI (in particular OPA) and thus averaging could be masking local effects of TMS within the ROIs. To avoid the issue, we performed a linear mixed effects analysis at the whole brain level (implemented in AFNI using 3dLME), which has the potential to reveal local effects of TMS if spatially consistent across participants as well as potential downstream effects that fall outside our *a priori* defined ROIs. These analyses were conducted on the surface data (in each hemisphere separately) of the stimulated hemisphere (see **Supplementary Figure S3** for the non-stimulated hemisphere results). Analogous to the ROI analysis, we looked for fixed effects of Session, TMS-condition, Stimulus, Run and their potential interactions, whilst modeling participant as a random effect using a random intercept (but see **Supplementary Figure S5** for whole-brain effects modelled with random slopes for each participant).

At the whole brain level, significant clusters were evident for each main effect **(see Supplementary Figures S4)**, but only the interactions of Session by TMS-condition and Session by Run were significant at the selected cluster corrected statistical threshold (*q* = 0.000042). The Session by Run interaction revealed only a very small cluster on the ventral surface, close to the posterior boundary of PPA, which reflects on average a larger reduction in response at this location across runs in the Post sessions.

In line with our ROI analyses, we focused on the Session by TMS-condition interaction as it demonstrates differences in TMS induced effects across sessions (Pre, Post). Significant interaction effects were observed as localised clusters that either overlapped with, or were in close proximity to, our group-based definitions for OPA, OFA, PPA, FFA and parts of V1 (**Figure 4A**). While no effects were observed in the group-based MPA, a small cluster was present just inferior to the group ROI. Of particular note, we observed spatially localized effects within the stimulated OPA, as well as posterior OFA, coupled with more spatially extended effects within and adjacent to PPA.

**Figure 4:**
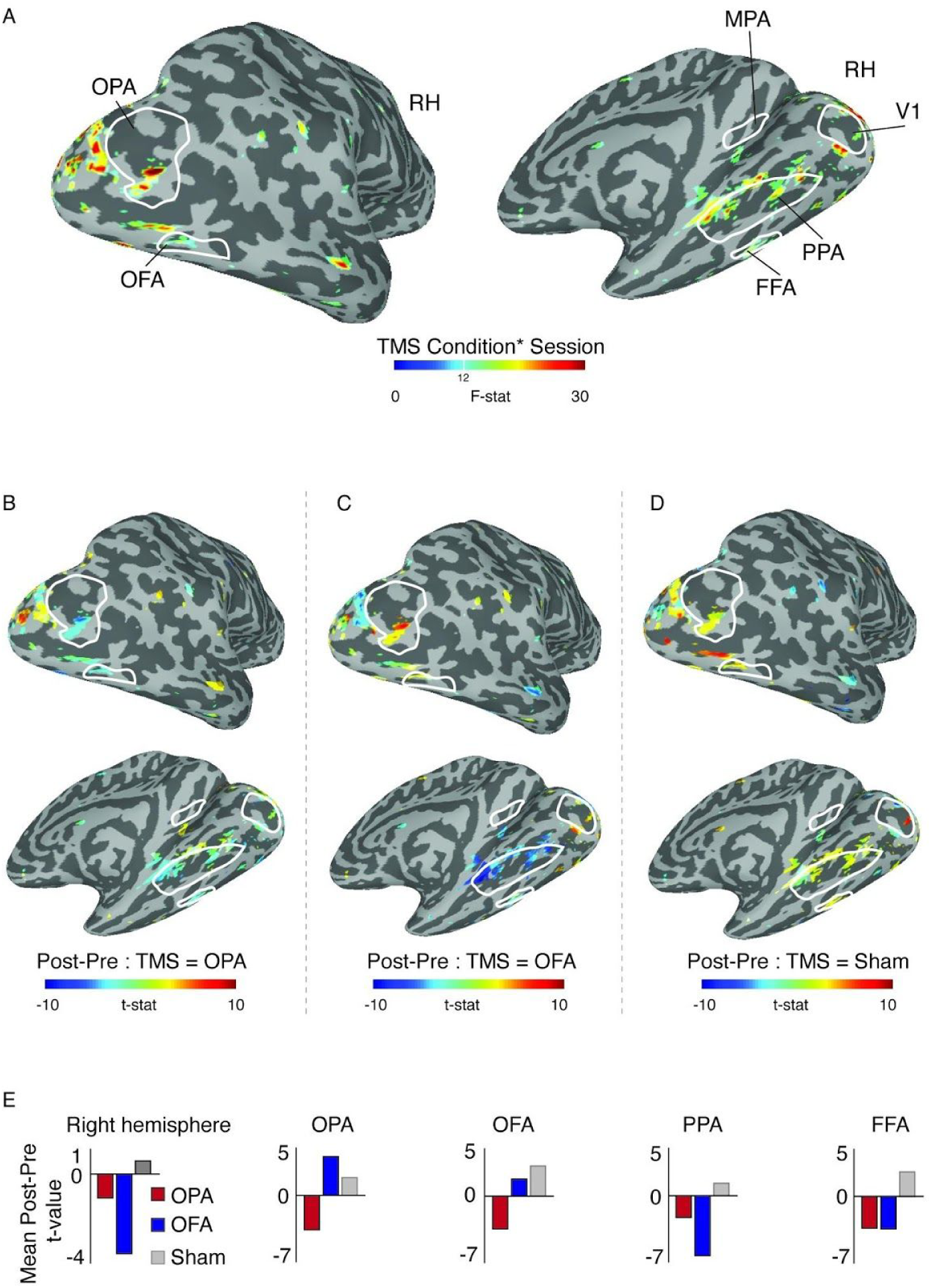
Linear mixed effects results in the stimulated right hemisphere. **A**, Locations of significant Session by TMS Condition interaction effects (p=0.0000024, q=0.000042) are overlaid onto lateral (top) and medial (bottom) views of a partially inflated surface reconstruction of the right hemisphere (sulci=dark gray, gyri=light gray). Interaction effects are evident overlapping with (or in close proximity to) group-based definitions of OPA, OFA (left), PPA, MPA, FFA(right) as well as early visual cortex. **B**, The effect of Session (Pre, Post) following OPA stimulation is overlaid onto the same views as A and thresholded on the interaction shown in A. Cold colors represent a decrease in response following TMS of OPA, with hot colors representing an increase. On the whole, TMS of OPA caused a reduction in response. Local decreases are evident inside OPA, OFA, anterior PPA and FFA. **C**, Same as B but for OFA stimulation. Here, positive responses are present on the lateral surface and within OPA and OFA specifically. On the ventral surface, responses are negative and numerically larger than those following OPA stimulation. **D**, Same as B but for sham condition. Responses are largely positive throughout the brain. **E**, Bars represent the mean t-value for the effect of Session during OPA (red), OFA (blue) and Sham (gray) conditions. Considering the entire hemisphere (left plot), both stimulation conditions resulted in reduced responses, whereas a positive response followed the Sham condition. Additional plots represent the mean difference in activation across the nodes showing a significant interaction effect within OPA, OFA, PPA and FFA. See **Supplementary Figure S5** for a version of this analysis that included random slopes for each participant in addition to random intercepts, yielding qualitatively similar results.

To understand what was driving the interaction in these clusters, we looked at the direction of the difference in activation across Sessions (Post - Pre), for each TMS condition separately. The group averaged maps are presented in **Figure 4 B-D** and the average difference within the entire hemisphere and in each ROI is presented in **Figure 4E**. In these figures, positive values represent an increase in response in the Post session (over Pre), whereas negative values represent a reduction in response in the Post session (over Pre). Note that these values are collapsed across the factors Stimulus and Run.

During Sham baseline, effects in each ROI were almost exclusively positive (albeit small in magnitude), reflecting on average an increased response during the Post session. In downstream ventral temporal regions PPA and FFA, consistent with the ROI analyses, the interaction appears to be driven by reduced responses as a result of either stimulating OPA or OFA. In the stimulated regions, the pattern is slightly different for each ROI. For clusters within both OPA and OFA there were decreased responses in a local region for OPA stimulation but increased responses for OFA stimulation. However, in OPA but not OFA these increases were greater than those for Sham.

Overall, the whole-brain analyses show that whilst TMS of both OPA and OFA elicited an overall reduction in response relative to the Sham baseline, the impact of stimulation also differed between stimulation sites. Relative to Sham, OPA stimulation led to a local decrease within OPA, OFA, PPA and FFA, whereas OFA stimulation led to a local increase in response in OPA, a small decrease in OFA, and a more pronounced reduction in downstream areas PPA and FFA.

### Behavioral data analysis

This study was not optimized to examine the behavioral impact of TBS stimulation. However, given the widespread nature of the observed TBS effects, we performed an exploratory analysis to test whether participant performance was affected by TBS, considering both repeat performance and reaction times (see **Supplementary Methods 2**). No behavioral effects of TBS were observed (see **Supplementary Results 4 and Supplementary Figure S6)**.

## Discussion

Our results show that theta-burst stimulation of a scene-selective region in the lateral occipital cortex (OPA) reduces BOLD responses in a scene-selective region in the ventral temporal cortex (PPA). Although we did not observe effects of TMS in OPA itself when conducting a ROI-based analysis that averaged activation values across the entire ROI, the whole-brain analysis did reveal the presence of a local effect within OPA as a result of stimulation. While we thus observed both local and downstream effects of TMS on fMRI responses in scene regions to visual stimuli, our findings did not fully confirm our predictions. Specifically, we predicted that TMS effects would reduce responses in downstream scene-selective regions in a stimulus- and stimulation-site specific manner. However, responses in PPA were reduced across all stimulus categories, and were similarly reduced when stimulating an active control region that was face-selective (OFA). Conversely, TMS-induced effects were observed in face-selective FFA when stimulating not only face-selective OFA but also when stimulating scene-selective OPA. Furthermore, only limited TMS-induced effects were observed in the proximity of a second downstream scene-selective region, MPA. Collectively, these results show that effects of theta-burst TMS to the lateral occipital cortex are not constrained to a single category-selective network of regions. Below we discuss the implications of these results further and highlight the importance of an active control site in studies such as this.

### Lack of stimulation site-specificity in downstream TMS effects

Our results clearly indicate that stimulation of both OPA and OFA causally decreased responses relative to the Sham baseline. The OFA stimulation served primarily as a control condition for our main research question investigating causal interactions between scene-selective regions OPA, PPA and MPA, and our experiment was designed to investigate scene processing. However, we also reported TMS induced effects in FFA, which showed that stimulating OPA in fact also reduced responses in FFA (similar to stimulating OFA). The fact that TMS of both OPA and OFA elicited downstream reductions in the same areas speaks to whether or not these scene- and face-selective networks can be separated cleanly with theta-burst stimulation and fMRI. Online TMS experiments (Pitcher et al., 2009; Dilks et al., 2013) have demonstrated that such separation is possible when repetitive TMS is paired with indices of behavioural performance (e.g. reaction time, accuracy), but such clear separation might be less likely with TBS and fMRI. Alternatively, this lack of separation following TBS may reflect the underlying state of the stimulated regions. For example, TMS has been shown to induce different effects depending on whether stimulation occurs during a ‘passive-phase’ as in the current experiment, or whether stimulation occurs whilst that region is actively engaged in a task (Silvanto et al., 2007). In the latter case, TMS has been shown to increase the activity of neural populations less engaged by the ongoing task demands.

Some evidence for separation between regions using TBS/fMRI has been shown within the face-network (Pitcher et al., 2014; 2017) using both static and dynamic face stimuli, but there the critical comparison was between stimulation of OFA and the posterior superior temporal sulcus (pSTS), which is farther away from OFA than OPA is and whose preference for dynamic stimuli could be seen as evidence for it belonging to a network separate from OFA (e.g. static v dynamic, Pitcher & Ungerleider, 2021). Prior TMS/fMRI work investigating cognitive control also highlighted the differential impact TMS can have following within-network versus between-network stimulation For example, Gratton and colleagues (2013) found widespread effects following TBS of the left anterior insula and left dorsolateral prefrontal cortex (DLPFC), but more focal effects when TBS was applied to primary somatosensory cortex (S1). The differential spread of TBS was interpreted as reflecting the wide-spread connections that the insula and DLPFC have to other regions involved in cognitive control as compared to S1’s. Placed in the context of the current study, the lack of site specificity could reflect OPA and OFA belonging to the same functional network in general despite their different categorical preferences.

A further possibility for the lack of site-specific effects in the current study is how the impact of TBS was operationalised. Here, our principal method for assessing the impact of TBS was a comparison of the fMRI responses before and after stimulation and not its impact on behavioural performance, which is often how TMS effects are demonstrated (Pitcher et al., 2007; Silson et al., 2013; Chen et al., 2016). We know of at least one prior TBS study that successfully dissociated between two close proximity sites in visual cortex using a behavioral measure (Chen et al., 2016). Unlike in our study, where the two-back task was not optimised for precise behavioral measurements, Chen and colleagues (2016) took advantage of carefully optimised psychophysical procedures that likely enabled the detection of a TBS effect.

We also assessed whether the lack of site-specific effects was due to the two target sites being in closer proximity to each other in certain participants over others. To look at this, we calculated the Euclidean distance between target sites in each participant (mean = 49.33mm, std = 11.96mm). For 15/16 participants, these distances were largely similar. Indeed, only a single participant had a distance > 1 std above the mean. Thus our results are unlikely to reflect closer proximity of target locations in a subset of participants.

Interestingly, TMS induced reductions in ventral temporal cortex appeared slightly stronger overall for stimulating OFA compared to stimulating OPA. It is possible that the strong impact of OFA stimulation on downstream areas of the ventral temporal cortex reflects its close proximity to VTC (compared with, for example, OPA). In all participants, OFA is located ventral and slightly anterior to OPA. Indeed, in many cases there is little cortex separating OFA on the lateral surface and FFA on the ventral surface. Any impact of OPA stimulation has much further to travel to affect VTC regions. Consistent with this suggestion, prior work (Rafique et al., 2017) reported stronger downstream effects following TMS of object-selective LO, as compared to scene-selective OPA, with LO located in close proximity to OFA on the lateral surface (Silson et al., 2016). Thus, the proximity differences with VTC between our stimulation sites could account for both the current and prior downstream VTC effects (Rafique et al., 2017).

### Lack of stimulus-specificity in TMS induced response

Both the ROI-based and whole-brain analysis did not indicate statistical evidence for stimulus-specificity of the TMS induced effects, suggesting that TMS similarly affected fMRI responses to all stimulus categories, rather than the specific category preferred by either the stimulated ROI or the downstream ROI. Prior evidence for stimulus-specific effects of TMS on fMRI responses is mixed. Stimulation of face-selective OFA has been found to reduce responses in downstream ventral temporal regions for both static faces and objects (Pitcher et al., 2014), but another study reported more extensive reductions for faces than for butterflies (Solomon-Harris et al., 2016). Stimulation of scene-selective OPA did not result in significant differences in reduced responses between scenes vs. objects in downstream PPA (Rafique et al., 2017), but stimulation of object-selective LO has been found to reduce responses to objects in LO but increase responses to scenes in PPA (Mullin et al., 2013). As we highlighted above however, behavioral effects induced by online TMS have consistently been found to be stimulus-specific. This suggests that, compared with neural interference induced by online TMS, consecutive TBS effects may lead to reduced fMRI BOLD responses that are more spread out across multiple category-processing networks. The behavioral impact of these more widespread response reductions is unclear. In the current study, we did not observe any behavioral impact, but as highlighted above, our experiment was not optimized to identify behavioral effects of the brain stimulation; this might for example be possible when using more tailored psychophysical methods (Chen et al., 2016). Future research is needed to ascertain the degree of stimulus-specificity of behavioral impact of consecutive TBS-fMRI on category-selective visual cortex.

### Lack of TMS-induced effects in MPA

Whilst TMS-induced effects were observed in downstream PPA, they were largely absent in MPA (although effects were proximal to MPA at the whole-brain level). There are two, not necessarily mutually exclusive, explanations for the lack of TMS-induced effect in MPA. First, it is possible that this lack of effect reflects the general weaker responses in MPA relative to the other ROIs (**Supplementary Figure 2)**. Indeed, positive responses in MPA were only elicited by half of the stimulus set (all scene categories). In contrast, our additional ROIs exhibited a much broader response profile whilst still maintaining category preferences. Given that our TMS effects were not stimulus specific, the overall weaker responses in MPA across our stimulus set could underpin the lack of effect observed. Second, the lack of effect could reflect the strength of MPAs connectivity with the other scene regions. Prior work from our group (Silson et al., 2016) and others (Baldassano et al., 2013) using resting-state functional connectivity demonstrated that MPA is more connected to PPA ventrally than OPA laterally. This lack of TMS induced effect in MPA could reflect its reduced connectivity with OPA.

### Importance of the active control site

Two types of controls were employed in order to test the specificity of TMS stimulation (compared to Sham), and the specificity of stimulating the scene-network (compared to a non-scene, but close proximity region, OFA). The inclusion of a Sham condition allowed us to test the effect of stimulation, but the active control site (OFA) allowed us to test whether any stimulation effects were specific to the scene-network. It is important to note that our interpretations would be dramatically different without the inclusion of this active control site. If we had only compared OPA stimulation with Sham, we would have reported that TBS to OPA leads to both a local decrease within OPA itself, as well as downstream decreases in PPA, and we could have interpreted this as evidence for a key role of OPA in routing visual information through the scene network. Such an interpretation however is not valid when the results from our active control are taken into account. Here, we saw that reductions in PPA were also evident when stimulating face-selective OFA, suggesting that causal effects in scene-selective ventral temporal cortex can also be elicited by stimulating visually responsive regions that do not share the same category preference. The same pattern was observed for reductions in face-selective ventral temporal cortex (FFA) when stimulating scene-selective lateral-occipital cortex. Whilst this makes the interpretation more challenging, it is important to reflect on the potentially misleading picture that would have been painted had we not included an active control condition. Recent TMS work, both behaviourally (Julien et al., 2016) and in combination with fMRI (Wang et al., 2014; Wang & Voss, 2015; Pitcher et al., 2017, Thakral et al., 2020) have compared only one-active site with either vertex stimulation (Julien et al., 2016; Thakral et al., 2020) or Sham (Wang et al., 2014; Wang & Voss, 2015), complicating site-specific interpretations of TMS results. For instance, comparing the impact of visual cortex and vertex stimulation on a visual discrimination task tells us that visual cortex is causally involved in visual perception, but not whether the targeted visual region is uniquely responsible for the visual discrimination performance. For site specificity, TMS effects at the target location should be ideally compared to stimulation of a close proximity active control site that itself is not hypothesized to be critically involved in the visual process of interest (Pitcher et al., 2009, Silson et al., 2013; Solomon-Harris et al., 2016). We argue here that the inclusion of an active control site should be carefully considered in future TMS studies.

### Comparison of TMS induced effects within OPA versus OFA

Our whole brain analysis revealed a highly localised TMS effect within OPA. In OFA, however, the interaction effect was weaker in magnitude and less localised to our group-based definition. We considered the possibility that this might simply reflect less variability in stimulation site for OPA across participants as compared to OFA. To address this, we assessed the pairwise Euclidean distances between stimulation sites across participants and found that there was significantly less variance in stimulation location across participants for OPA compared to OFA (*one-tailed t-test: t(15) = 1*.*96, p*<0.03; **Supplementary Figure S7** and **Figure 1D**). It is thus possible that the increased variability in OFA stimulation site across subjects underpins the lower degree of overlap between our group-based OFA definition and the interaction effects revealed by our whole-brain linear mixed-effects analysis.

Another potential account for the larger local effect observed in OPA compared to OFA concerns OFAs proximity to brain locations characterized by fMRI signal distortion due to large draining veins. Of particular relevance here is the so-called *venous eclipse*, the lateral projection of the dural sinus which in certain individuals runs in close proximity to area V4, causing substantial distortions in the fMRI signal therein (Winawer et al., 2010). Our prior work on OFA (Silson et al., 2016) revealed that the ventral boundary of OFA is in close proximity to the same artefact. Here, we considered whether on average the proximity of OFA to the venous eclipse might spatially distort the signals we measure from this region. At the group level, the ventral boundary of OFA indeed abuts the lateral projection of the dural sinus (**Supplementary Figure S8**). It is possible therefore that the dural sinus distorted the signals in and around the OFA resulting in reduced overlap with our group-based ROI definition.

Alternatively, it is possible that inaccuracies in ROI alignment between sessions could lead to responses being shifted away from our original OFA definitions. To address this possibility we conducted several complementary analyses, which we believe rule out this possibility and provide confidence in our analysis approach. First, we examined the spatial overlap between our original ROI definitions (taken from the independent localiser) and those computed from each of the four pre TMS runs. Despite being computed on individual runs as opposed to the average of six, all pre TMS ROIs fell within the boundaries of our original ROI definitions, although considerably smaller in areal extent (**Supplementary Figure S1D**). Second, we calculated the Euclidean distance between the centre of mass coordinate of the original ROI (from the independent localiser session) and the centre of mass coordinate from an ROI defined from each of the four Sham Pre runs, before averaging these values across runs. Notwithstanding a single outlier, the average centre of mass distance was largely equivalent between the two target sites. Indeed, a paired t-test showed no significant difference in centre of mass shift between the two sites (t(15)=0.91, p=0.37). Third, we calculated the mean Euclidean distance between the target voxel in each ROI and the peak voxel from the equivalent contrast in each of the four sham pre runs, before averaging these values across runs. On average, displacements were ∼10mm for both target sites (**Supplementary Figure S1A)**. Indeed, a paired t-test showed no significant difference in shift distance (t(15)=0.08, p=0.93). Taken together, we are confident that the lack of local effect in OFA is not due to inaccuracies in localisation or alignment across sessions.

### TBS versus other types of TMS

The choice of TMS protocol can impact the spatial extent (Barker et al., 1985; Walsh & Cowey, 2000), temporal window and magnitude of TMS-induced effects (Allen et al., 2007). These factors also interact with how one operationalises the impact of TMS, such as whether one is measuring neuronal firing (Allen et al., 2007), phosphene generation (Walsh & Cowey, 2000), reaction times and discrimination accuracy (Pitcher et al., 2007; Dilks et al., 2013) or fMRI responses (Pitcher et al., 2014; Wang et al., 2014; Rafique et al., 2017). Here, we employed a TBS paradigm, which enabled us to both minimize the time between Pre and Post sessions (60s TBS + ∼2mins set-up) and potentially capture the prolonged impact of TBS stimulation on neuronal responses (Huang et al., 2005; Pitcher et al., 2014; 2017). However, despite the advantages of TBS, it is possible that more focal effects could have been elicited with another paradigm, such as repetitive TMS (Mullin & Steeves, 2013; Rafique et al., 2017) or single-pulse TMS, both of which can be delivered safely at higher intensities than TBS and have been used previously to distinguish between the functional roles of close proximity targets (Pitcher et al., 2009; Dilks et al., 2013). Importantly, this is just one explanation for the pattern of results we observed. As highlighted above, it is also possible that the current data better reflect either the different effects of stimulation during active or passive states (Silvanto et al., 2007), or the different ways one can operationalize the impact of TMS.

### Decreased responses across runs

Although it is common practice in fMRI research to have participants perform the same tasks in multiple runs, it is not without its drawbacks. Prior work (Meshulam & Malach, 2016) demonstrated a systematic decrease in fMRI response during repeated runs of a face gender/age discrimination task. The reduction in response magnitude across participants was correlated with subjective ratings of increased boredom and decreased attention on task. Consistent with this prior work (Meshulam & Malach, 2016), we also observed systematic reductions in response magnitude as a function of run in our ROI analyses, coupled with a decrease in behavioral accuracy across runs (**Supplementary Figure S6B**), that were largely equivalent in both Pre and Post TBS sessions. At the whole brain level, significant response reductions were present throughout much of the visual cortex (**Supplementary Figure S4D**). It is therefore possible that our repeating task structure and the subsequent effect of run masked some of the effects of TMS. The prominent effect of run reported here and in prior work (Meshulam & Malach, 2016) is an important consideration when designing fMRI tasks and concurrent TMS/fMRI experiments in particular.

## Conclusion

Local effects of TBS stimulation to scene- and face-selective cortex were observed within the stimulated regions, but these were insensitive to the preferred category of the stimulated region. Downstream effects were observed to be focal to scene and face-selective regions, but these effects were not constrained by the preferred category of the stimulated region, since similar TMS effects were observed when stimulating either a scene- or a face-selective lateral occipital region. Collectively, these results show that effects of theta-burst TMS to the lateral occipital cortex are not constrained to a single category-selective network, suggesting that information processing in different groups of category-selective regions (e.g. scene, face) may be less independent that one might assume based on their distinct category preferences alone. It is vital however to also recognise that the evidence for the independence of these different category-selective networks changes with experimental decisions, such as the type of TMS and how such TMS effects are operationalised. Finally, our results highlight the importance of a close proximity control stimulation site to fully interpret the site-specificity of TMS effects.

## Acknowledgements

This work was supported by the Intramural Research Program of the National Institute of Mental Health (ZIAMH002909), a Rubicon Fellowship from the Netherlands Organization for Scientific Research (NWO) to IIAG, and a BBSRC (UK) project grant (BB/P006981/1) to DP. Data were collected under National Institute of Mental Health Clinical Study Protocols 93-M-0170 (NCT00001360) and 12-M-0128 (NCT01617408).

