## Supplementary Materials for "Theta-burst TMS of lateral occipital cortex reduces BOLD responses across category-selective areas in ventral temporal cortex"

#### Table of contents

**Supplementary Methods** **2**

1. TMS-fMRI session counterbalancing scheme 2
2. Behavioral data analysis 2

**Supplementary Results** **3**

1. ROI analysis: full statistical tables for omnibus ANOVAs 3
    - 1.1 ANOVA results for scene-selective OPA and PPA 3
    - 1.2 ANOVA results for face-selective OFA and FFA 4
    - 1.3 ANOVA results for other ROIs: MPA and V1 5
2. ROI analysis: linear mixed effect models 6
    - 2.1 Linear mixed effects model comparison 7
        - 2.1.1 Random effects: R lmer formulas for model comparison 7
        - 2.1.2 Random effects: Likelihood ratio tests 8
        - 2.1.3 Fixed effects: R lmer formulas for model comparison 8
        - 2.1.4 Fixed effects: Likelihood ratio tests 9
    - 2.2 ANOVA results for model with random slopes 10
        - 2.2.1 OPA and PPA 10
        - 2.2.2 FFA and OFA 11
        - 2.2.3 MPA and V1 12
3. Whole brain analysis: additional linear mixed effects results 13
4. Behavioral data analysis: linear mixed effects results 14

**Supplementary Figures** **16**

- S1: Target and localizer ROI accuracy 16
- S2: ROI analysis for MPA and V1 17
- S3: Whole brain linear mixed effects in the left hemisphere 18
- S4: Whole brain main effects 19
- S5: Whole brain linear mixed effects with random slopes 20
- S6: Behavioral data (d’ and reaction times) 21
- S7: Variability in stimulation sites across participants 22
- S8: Proximity of OFA and the venous eclipse 23

**Supplementary References** **24**

### Supplementary Methods

#### 1. TMS-fMRI session counterbalancing scheme

| | stimulation | | | |
|---|---|---|---|---|
| **subject #** | OPA | OFA | Sham | run order |
| **1** | 1 | 2 | 3 | 2301 |
| **2** | 1 | 2 | 3 | 3012 |
| **3** | 1 | 2 | 3 | 0123 |
| **4** | 1 | 2 | 3 | 3021 |
| **5** | 2 | 1 | 3 | 1230 |
| **6** | 2 | 1 | 3 | 1032 |
| **7** | 2 | 1 | 3 | 2103 |
| **8** | 2 | 1 | 3 | 3210 |
| **9** | 3 | 2 | 1 | 0321 |
| **10** | 3 | 2 | 1 | 2301 |
| **11** | 3 | 2 | 1 | 3012 |
| **12** | 3 | 2 | 1 | 0123 |
| **13** | 2 | 3 | 1 | 1230 |
| **14** | 2 | 3 | 1 | 1032 |
| **15** | 2 | 3 | 1 | 2103 |
| **16** | 2 | 3 | 1 | 3210 |

#### 2. Behavioral data analysis

The recorded behavioral responses were used to calculate discriminability measure d' (MacMillan & Creelman, 1991). In this analysis, correct responses to image repeats (occurring between 1-3 times for each stimulus block) were scored as Hits, while button presses to non-repeats were scored as False Alarms. Responses were marked as correct if the button press occurred after the onset of the repeated image before the onset of the third following image presentation (resulting in a 2400ms response window). Responses made outside that window but during stimulus presentation were marked as incorrect. Reaction times (RTs) were calculated based on the correct trials and reflected the time elapsed between the button press and the onset of the repeated image. Discriminability values and RTs were calculated separately for each subject, category and run. Scores of 1.0 or 0.0 for hits and false alarms for a given category within a run were subjected to a standard correction whereby half a trial was either subtracted or added to the actual score. The resulting d' and RT measures were fitted with a linear mixed effects model incorporating the same factors as the main fMRI analyses

(Fixed effects: TMS condition, Session, Run and Stimulus; random effects: subject). The behavioral data table and R analysis script are provided on the OSF (https://osf.io/6nq7r/).

### Supplementary Results

#### 1. ROI analysis: full statistical tables for omnibus ANOVAs

The tables show the statistical output for the omnibus ANOVAs that were run on the fitted linear mixed effects models to identify potential effects of TBS in OPA and PPA (**section 1.1**; associated data in **Figure 2**), OFA and FFA (**section 1.2**; **Figure 3**) and MPA and V1 (**section 1.3**; **Figure S1**).

##### 1.1 ANOVA results for scene-selective OPA and PPA

| Type III Analysis of Variance Table with Satterthwaite's method | | | | | | | |
|---|---|---|---|---|---|---|---|
| | Sum Sq | Mean Sq | NumDF | DenDF | F value | Pr(>F) | Pr(>F) bonf |
| ***OPA*** | | | | | | | |
| TMScondition | 0.00 | 0.00 | 2 | 2857 | 0.02 | 0.979 | |
| Session | 0.01 | 0.01 | 1 | 2857 | 0.07 | 0.796 | |
| **Stimulus** | **246.06** | **35.15** | **7** | **2857** | **370.99** | **0.000** | **0.000** |
| **Run** | **21.76** | **7.25** | **3** | **2857** | **76.56** | **0.000** | **0.000** |
| TMScondition:Session | 0.18 | 0.09 | 2 | 2857 | 0.93 | 0.395 | |
| TMScondition:Stimulus | 1.00 | 0.07 | 14 | 2857 | 0.75 | 0.719 | |
| Session:Stimulus | 0.48 | 0.07 | 7 | 2857 | 0.72 | 0.657 | |
| TMScondition:Run | 0.93 | 0.16 | 6 | 2857 | 1.64 | 0.133 | |
| Session:Run | 0.98 | 0.33 | 3 | 2857 | 3.44 | 0.016 | 0.097 |
| Stimulus:Run | 1.14 | 0.05 | 21 | 2857 | 0.57 | 0.938 | |
| TMScondition:Session: Stimulus | 0.81 | 0.06 | 14 | 2857 | 0.61 | 0.860 | |
| TMScondition:Session:Run | 0.49 | 0.08 | 6 | 2857 | 0.87 | 0.518 | |
| TMSconditionStimulus:Run | 1.50 | 0.04 | 42 | 2857 | 0.38 | 1.000 | |
| Session:Stimulus:Run | 1.40 | 0.07 | 21 | 2857 | 0.70 | 0.835 | |
| TMScondition:Session: Stimulus:Run | 1.29 | 0.03 | 42 | 2857 | 0.33 | 1.000 | |
| ***PPA*** | | | | | | | |
| **TMScondition** | **0.85** | **0.42** | **2** | **2857** | **5.48** | **0.004** | **0.025** |
| **Session** | **2.46** | **2.46** | **1** | **2857** | **31.93** | **0.000** | **0.000** |
| **Stimulus** | **266.31** | **38.04** | **7** | **2857** | **493.93** | **0.000** | **0.000** |
| **Run** | **26.63** | **8.88** | **3** | **2857** | **115.25** | **0.000** | **0.000** |
| **TMScondition:Session** | **1.41** | **0.70** | **2** | **2857** | **9.14** | **0.000** | **0.001** |

| | | | | | | | |
|---|---|---|---|---|---|---|---|
| TMScondition:Stimulus | 1.19 | 0.09 | 14 | 2857 | 1.10 | 0.348 | |
| Session:Stimulus | 0.12 | 0.02 | 7 | 2857 | 0.23 | 0.979 | |
| TMScondition:Run | 0.60 | 0.10 | 6 | 2857 | 1.30 | 0.255 | |
| Session:Run | 0.96 | 0.32 | 3 | 2857 | 4.14 | 0.006 | 0.037 |
| Stimulus:Run | 1.79 | 0.09 | 21 | 2857 | 1.11 | 0.333 | |
| TMScondition:Session: Stimulus | 0.28 | 0.02 | 14 | 2857 | 0.26 | 0.997 | |
| TMScondition:Session:Run | 0.43 | 0.07 | 6 | 2857 | 0.94 | 0.467 | |
| TMScondition: Stimulus:Run | 1.44 | 0.03 | 42 | 2857 | 0.44 | 0.999 | |
| Session:Stimulus:Run | 1.05 | 0.05 | 21 | 2857 | 0.65 | 0.884 | |
| TMScondition:Session: Stimulus:Run | 1.38 | 0.03 | 42 | 2857 | 0.43 | 1.000 | |

1.2 ANOVA results for face-selective OFA and FFA

| Type III Analysis of Variance Table with Satterthwaite's method | | | | | | | |
|---|---|---|---|---|---|---|---|
| | Sum Sq | Mean Sq | NumDF | DenDF | F value | Pr(>F) | Pr(>F) bonf |
| ***OFA*** | | | | | | | |
| **TMScondition** | **1.33** | **0.67** | **2** | **2857** | **5.92** | **0.003** | **0.016** |
| Session | 0.45 | 0.45 | 1 | 2857 | 4.01 | 0.045 | 0.272 |
| **Stimulus** | **121.17** | **17.31** | **7** | **2857** | **153.77** | **0.000** | **0.000** |
| **Run** | **32.79** | **10.93** | **3** | **2857** | **97.09** | **0.000** | **0.000** |
| TMScondition:Session | 0.12 | 0.06 | 2 | 2857 | 0.51 | 0.598 | |
| TMScondition:Stimulus | 1.58 | 0.11 | 14 | 2857 | 1.00 | 0.445 | |
| Session:Stimulus | 0.36 | 0.05 | 7 | 2857 | 0.46 | 0.865 | |
| TMScondition:Run | 0.59 | 0.10 | 6 | 2857 | 0.88 | 0.509 | |
| **Session:Run** | **2.07** | **0.69** | **3** | **2857** | **6.14** | **0.000** | **0.002** |
| Stimulus:Run | 2.43 | 0.12 | 21 | 2857 | 1.03 | 0.426 | |
| TMScondition:Session: Stimulus | 0.68 | 0.05 | 14 | 2857 | 0.43 | 0.964 | |
| TMScondition:Session:Run | 0.87 | 0.15 | 6 | 2857 | 1.29 | 0.258 | |
| TMScondition:Stimulus:Run | 2.52 | 0.06 | 42 | 2857 | 0.53 | 0.994 | |
| Session:Stimulus:Run | 1.56 | 0.07 | 21 | 2857 | 0.66 | 0.875 | |
| TMScondition:Session: Stimulus:Run | 2.08 | 0.05 | 42 | 2857 | 0.44 | 0.999 | |
| ***FFA*** | | | | | | | |
| **TMScondition** | **2.60** | **1.30** | **2** | **2857** | **7.84** | **0.000** | **0.002** |

| | | | | | | | |
|---|---|---|---|---|---|---|---|
| Session | 0.69 | 0.69 | 1 | 2857 | 4.18 | 0.041 | 0.246 |
| **Stimulus** | **192.97** | **27.57** | **7** | **2857** | **166.46** | **0.000** | **0.000** |
| **Run** | **32.87** | **10.96** | **3** | **2857** | **66.16** | **0.000** | **0.000** |
| **TMScondition:Session** | **3.75** | **1.88** | **2** | **2857** | **11.33** | **0.000** | **0.000** |
| TMScondition:Stimulus | 2.33 | 0.17 | 14 | 2857 | 1.00 | 0.445 | |
| Session:Stimulus | 0.79 | 0.11 | 7 | 2857 | 0.68 | 0.692 | |
| TMScondition:Run | 0.72 | 0.12 | 6 | 2857 | 0.73 | 0.627 | |
| **Session:Run** | **2.99** | **1.00** | **3** | **2857** | **6.02** | **0.000** | **0.003** |
| Stimulus:Run | 3.03 | 0.14 | 21 | 2857 | 0.87 | 0.629 | |
| TMScondition:Session: Stimulus | 0.58 | 0.04 | 14 | 2857 | 0.25 | 0.998 | |
| TMScondition:Session:Run | 0.41 | 0.07 | 6 | 2857 | 0.41 | 0.872 | |
| TMScondition:Stimulus:Run | 3.26 | 0.08 | 42 | 2857 | 0.47 | 0.999 | |
| Session:Stimulus:Run | 2.74 | 0.13 | 21 | 2857 | 0.79 | 0.739 | |
| TMScondition:Session: Stimulus:Run | 2.44 | 0.06 | 42 | 2857 | 0.35 | 1.000 | |

1.3 ANOVA results for other ROIs: MPA and V1

| Type III Analysis of Variance Table with Satterthwaite's method | | | | | | | |
|---|---|---|---|---|---|---|---|
| | Sum Sq | Mean Sq | NumDF | DenDF | F value | Pr(>F) | Pr(>F) bonf |
| ***MPA*** | | | | | | | |
| TMScondition | 0.27 | 0.14 | 2 | 2857 | 1.46 | 0.233 | |
| **Session** | **2.44** | **2.43** | **1** | **2857** | **26.23** | **0.000** | **0.000** |
| **Stimulus** | **186.88** | **26.70** | **7** | **2857** | **287.64** | **0.000** | **0.000** |
| **Run** | **11.80** | **3.93** | **3** | **2857** | **42.37** | **0.000** | **0.000** |
| TMScondition:Session | 0.20 | 0.10 | 2 | 2857 | 1.07 | 0.345 | |
| TMScondition:Stimulus | 1.60 | 0.11 | 14 | 2857 | 1.23 | 0.247 | |
| Session:Stimulus | 0.27 | 0.04 | 7 | 2857 | 0.41 | 0.895 | |
| TMScondition:Run | 0.97 | 0.16 | 6 | 2857 | 1.74 | 0.109 | |
| Session:Run | 0.22 | 0.07 | 3 | 2857 | 0.78 | 0.507 | |
| Stimulus:Run | 2.09 | 0.10 | 21 | 2857 | 1.07 | 0.372 | |
| TMScondition:Session: Stimulus | 0.90 | 0.06 | 14 | 2857 | 0.69 | 0.785 | |
| TMScondition:Session:Run | 0.88 | 0.15 | 6 | 2857 | 1.58 | 0.150 | |
| TMScondition:Stimulus:Run | 2.00 | 0.05 | 42 | 2857 | 0.51 | 0.996 | |
| Session:Stimulus:Run | 1.54 | 0.07 | 21 | 2857 | 0.79 | 0.734 | |
| TMScondition:Session: Stimulus:Run | 2.64 | 0.06 | 42 | 2857 | 0.68 | 0.944 | |

| | | | | | | | |
|---|---|---|---|---|---|---|---|
| ***V1*** | | | | | | | |
| **TMScondition** | **1.32** | **0.66** | **2** | **2857** | **4.88** | **0.008** | **0.046** |
| Session | 0.00 | 0.00 | 1 | 2857 | 0.01 | 0.913 | |
| **Stimulus** | **32.62** | **4.66** | **7** | **2857** | **34.37** | **0.000** | **0.000** |
| **Run** | **16.74** | **5.58** | **3** | **2857** | **41.15** | **0.000** | **0.000** |
| TMScondition:Session | 0.36 | 0.18 | 2 | 2857 | 1.32 | 0.267 | |
| TMScondition:Stimulus | 0.91 | 0.07 | 14 | 2857 | 0.48 | 0.945 | |
| Session:Stimulus | 0.27 | 0.04 | 7 | 2857 | 0.28 | 0.962 | |
| TMScondition:Run | 0.97 | 0.16 | 6 | 2857 | 1.20 | 0.305 | |
| Session:Run | 1.59 | 0.53 | 3 | 2857 | 3.91 | 0.008 | 0.051 |
| Stimulus:Run | 2.87 | 0.14 | 21 | 2857 | 1.01 | 0.447 | |
| TMScondition:Session: Stimulus | 1.21 | 0.09 | 14 | 2857 | 0.64 | 0.835 | |
| TMScondition:Session:Run | 0.46 | 0.08 | 6 | 2857 | 0.56 | 0.761 | |
| TMScondition:Stimulus:Run | 2.75 | 0.07 | 42 | 2857 | 0.48 | 0.998 | |
| Session:Stimulus:Run | 2.11 | 0.10 | 21 | 2857 | 0.74 | 0.793 | |
| TMScondition:Session: Stimulus:Run | 2.19 | 0.05 | 42 | 2857 | 0.39 | 1.000 | |

#### 2. ROI analysis: linear mixed effect models

Our linear mixed effects model specification included a random intercept for participants, allowing for the statistical model to take into account the high inter-participant variability in overall fMRI response. While less often applied in neuroimaging research, mixed effects models can incorporate more complex random effects structure, including random slopes for each within-subject factor and their interactions, allowing the model to take into account variability across participants from session to session, for example, or variability in participant's sensitivity to the experimental manipulation. We explored the inclusion of these factors in our ROI analyses using a model selection procedure (Meteyard & Davies, 2020; Brown 2020), whereby increasingly complex random effects structures were fitted first followed by the inclusion of the fixed effects, and model selection was evaluated using likelihood ratio tests (LRTs; see section 2.1. The results suggested that the data would be better fit (higher likelihood) by a model that additionally included random slopes for Session and TMS-condition, but not their interaction, for a all of the ROIs (see **section 2.1** below for a full report of the model selection and associated tests, and OSF page https://osf.io/6nq7r for the R script used for the model comparison).

Importantly, however, the overall pattern of results was qualitatively identical to that reported in the simpler, random intercept-only model, indicating TBS-induced response reductions in PPA and FFA but not OPA, OFA, MPA or V1 (see **section 2.2** below). A whole brain 3dLME analysis with the random effects structure that best fitted the ROI data also showed a qualitatively similar pattern to the main analysis (see **section 3** and **Figure S8**).

#### 2.1 Linear mixed effects model comparison

Following recommendations outlined in Meteyard & Davies (2020), we started with fitting random effects only (no fixed effects) to identify the best-fitting random effects structure. The respective model specifications are presented in **section 2.1.1**. The first model includes only a random intercept for subject; the second and third models add random slopes for TMScondition (OPA, OFA, Sham) and Session (Pre vs. Post), respectively. Additional models that included the interaction for Session and TMScondition or additional random effects resulted in singular or non converging models for all of the ROIs, and are therefore marked in gray. LRT results for each ROI comparing models 1,2 and 3 are presented in **section 2.1.2**. At the next step, we added fixed effects while keeping the random effects structure constant. The reference model (Model 1) contained no fixed effects, while model 2 included the manipulated within-subject factors. Finally, model 3 additionally included a fixed effect of Run. Model 3 performed resulted in the best fits in all ROIs. The model specifications and LRT results for each ROI are presented in **sections 2.1.3** and **2.1.4**, respectively.

##### *2.1.1 Random effects: R lmer formulas for model comparison*

| **Model name** | **Description** | **Formula** |
|---|---|---|
| Model R0 | random intercept only | lmer(fMRIresponse ~ 1 + (1\|subject) |
| Model R1 | random intercept and slope for TMScondition | lmer(fMRIresponse ~ 1 + (TMScondition\|subject) |
| Model R2 | random intercept and slope for TMScondition and Session | lmer(fMRIresponse ~ 1 + (TMScondition+Session\|subject) |
| Model R3 | random intercept and slope for TMS condition and Session interaction | lmer(fMRIresponse ~ 1 + (TMScondition*Session\|subject) |
| Model R4 | random intercept and slope for TMScondition and Session and Run | lmer(fMRIresponse ~ 1 + (TMScondition+Session+Run\|subject) |
| Model R5 | random intercept and slope for TMScondition and Session and Stimulus | lmer(fMRIresponse ~ 1 + (TMScondition+Session+Stimulus\|subject) |
| Model R6 | random intercept and slopes for all within-subject factors | lmer(fMRIresponse ~ 1 + (TMScondition+Session+Run+Stimulus\|subject) |

*2.1.2 Random effects: Likelihood ratio tests*

| | Df | AIC | BIC | logLik | deviance | Chisq | Df | Pr(>Chisq) |
|---|---|---|---|---|---|---|---|---|
| ***OPA*** | | | | | | | | |
| Model R0 | 3 | 3514.2 | 3532.3 | -1754.1 | 3508.2 | | | |
| Model R1 | 8 | 3456.7 | 3504.9 | -1720.3 | 3440.7 | 67.5 | 5 | **3.4E-13** |
| Model R2 | 12 | 3444.6 | 3516.9 | -1710.3 | 3420.6 | 20.1 | 4 | **4.8E-04** |
| ***PPA*** | | | | | | | | |
| Model R0 | 3 | 3401.4 | 3419.5 | -1697.7 | 3395.4 | | | |
| Model R1 | 8 | 3331 | 3379.2 | -1657.5 | 3315.0 | 80.4 | 5 | **7.0E-16** |
| Model R2 | 12 | 3316.8 | 3389.1 | -1646.4 | 3292.8 | 22.2 | 4 | **1.8E-04** |
| ***OFA*** | | | | | | | | |
| Model R0 | 3 | 3187.9 | 3206 | -1591 | 3181.9 | | | |
| Model R1 | 8 | 3111 | 3159.2 | -1547.5 | 3095.0 | 86.9 | 5 | **2.2E-16** |
| Model R2 | 12 | 3093.2 | 3165.5 | -1534.6 | 3069.2 | 25.8 | 4 | **3.5E-05** |
| ***FFA*** | | | | | | | | |
| Model R0 | 3 | 4385.4 | 4403.5 | -2189.7 | 4379.4 | | | |
| Model R1 | 8 | 4124.2 | 4172.4 | -2054.1 | 4108.2 | 271.2 | 5 | **2.20E-16** |
| Model R2 | 12 | 4105.9 | 4178.2 | -2041 | 4081.9 | 26.3 | 4 | **2.75E-05** |
| ***MPA*** | | | | | | | | |
| Model R0 | 3 | 3102.9 | 3121 | -1548.5 | 3096.9 | | | |
| Model R1 | 8 | 3080.1 | 3128.3 | -1532.1 | 3064.1 | 32.778 | 5 | **4.17E-06** |
| Model R2 | 12 | 3071.9 | 3144.2 | -1523.9 | 3047.9 | 16.26 | 4 | **0.002689** |
| ***V1*** | | | | | | | | |
| Model R0 | 3 | 2965.2 | 2983.3 | -1479.6 | 2959.2 | | | |
| Model R1 | 8 | 2770.6 | 2818.8 | -1377.3 | 2754.6 | 204.6 | 5 | **2.20E-16** |
| Model R2 | 12 | 2685.3 | 2757.7 | -1330.7 | 2661.3 | 93.237 | 4 | **2.20E-16** |

*2.1.3 Fixed effects: R lmer formulas for model comparison*

| **Model name** | **Description** | **Formula** |
|---|---|---|
| Model F0 | no fixed effects w/ best fitting random effects (model R2;see 2.1.2) * | lmer(fMRIresponse ~ 1 + (TMScondition+Session\|subject) |

| | | |
|---|---|---|
| Model F1 | fixed effects for experimentally manipulated within-subject factors | lmer(fMRIresponse ~ 1 + Stimulus*Session*TMScondition + (TMScondition+Session\|subject) |
| Model F2 | fixed effects for experimentally manipulated within-subject factors + Run | lmer(fMRIresponse ~ 1 + Stimulus*Session*TMScondition*Run + (TMScondition+Session\|subject) |

* Models R2 and F0 have identical model formulas, but their estimated coefficients are not numerically identical, because the models to compare fixed effects were fitted with maximum likelihood estimation (ML) rather than restricted maximum likelihood (REML) (see Brown, 2020; Meteyard and Davies, 2020).

*2.1.4 Fixed effects: Likelihood ratio tests*

| | Df | AIC | BIC | logLik | deviance | Chisq | Df | Pr(>Chisq) |
|---|---|---|---|---|---|---|---|---|
| ***OPA*** | | | | | | | | |
| Model F0 | 12 | 3440.5 | 3512.9 | -1708.3 | 3416.5 | | | |
| Model F1 | 59 | 1550.5 | 1906.1 | -716.3 | 1432.5 | 1984 | 47 | **2.20E-16** |
| Model F2 | 203 | 1481 | 2704.6 | -537.5 | 1075 | 357.52 | 144 | **2.20E-16** |
| ***PPA*** | | | | | | | | |
| Model F0 | 12 | 3313.3 | 3385.6 | -1644.6 | 3289.3 | | | |
| Model F1 | 59 | 1017.3 | 1372.9 | -449.6 | 899.3 | 2390 | 47 | **2.20E-16** |
| Model F2 | 203 | 796.57 | 2020.2 | -195.3 | 390.57 | 508.69 | 144 | **2.20E-16** |
| ***OFA*** | | | | | | | | |
| Model F0 | 12 | 3089.2 | 3161.6 | -1532.6 | 3065.2 | | | |
| Model F1 | 59 | 2229.3 | 2584.9 | -1055.7 | 2111.3 | 953.95 | 47 | **2.20E-16** |
| Model F2 | 203 | 2083.5 | 3307.1 | -838.8 | 1677.5 | 433.76 | 144 | **2.20E-16** |
| ***FFA*** | | | | | | | | |
| Model F0 | 12 | 4103.2 | 4175.6 | -2039.6 | 4079.2 | | | |
| Model F1 | 59 | 3048.3 | 3403.9 | -1465.2 | 2930.3 | 1148.9 | 47 | **2.20E-16** |
| Model F2 | 203 | 2976.4 | 4200.0 | -1285.2 | 2570.4 | 359.86 | 144 | **2.20E-16** |
| ***MPA*** | | | | | | | | |
| Model F0 | 12 | 3067.6 | 3139.9 | -1521.8 | 3043.6 | | | |
| Model F1 | 59 | 1543.6 | 1899.2 | -712.8 | 1425.6 | 1618 | 47 | **2.20E-16** |
| Model F2 | 203 | 1571.4 | 2795.0 | -582.7 | 1165.4 | 260.19 | 144 | **1.04E-08** |
| ***V1*** | | | | | | | | |
| Model F0 | 12 | 2683.3 | 2755.7 | -1329.7 | 2659.3 | | | |
| Model F1 | 59 | 2492.9 | 2848.5 | -1187.4 | 2374.9 | 284.46 | 47 | **2.20E-16** |
| Model F2 | 203 | 2519.8 | 3743.4 | -1056.9 | 2113.8 | 261.07 | 144 | **8.49E-09** |

#### 2.2 ANOVA results for model with random slopes

This section contains the full omnibus ANOVA statistical results for the best-fitting alternative linear mixed effects model that includes all four fixed effects plus random slopes for TMScondition and Session for ROIs OPA and PPA **(2.2.1)**, FFA and OFA ( **2.2.2**) and MPA and V1 **(2.2.3)**.

##### *2.2.1 OPA and PPA*

| Type III Analysis of Variance Table with Satterthwaite's method | | | | | | | |
|---|---|---|---|---|---|---|---|
| | Sum Sq | Mean Sq | NumDF | DenDF | F value | Pr(>F) | Pr(>F) bonf |
| ***OPA*** | | | | | | | |
| TMScondition | 0.00 | 0.00 | 2 | 15 | 0.01 | 0.992 | |
| Session | 0.00 | 0.00 | 1 | 15 | 0.00 | 0.957 | |
| **Stimulus** | **246.06** | **35.15** | **7** | **2812** | **420.00** | **0.000** | **0.000** |
| **Run** | **22.08** | **7.36** | **3** | **2812** | **87.95** | **0.000** | **0.000** |
| TMScondition:Session | 0.17 | 0.08 | 2 | 2812 | 0.98 | 0.374 | |
| TMScondition:Stimulus | 1.00 | 0.07 | 14 | 2812 | 0.85 | 0.609 | |
| Session:Stimulus | 0.48 | 0.07 | 7 | 2812 | 0.81 | 0.577 | |
| TMScondition:Run | 0.95 | 0.16 | 6 | 2812 | 1.90 | 0.078 | . |
| Session:Run | 0.89 | 0.30 | 3 | 2812 | 3.54 | 0.014 | 0.084 |
| Stimulus:Run | 1.14 | 0.05 | 21 | 2812 | 0.65 | 0.884 | |
| TMScondition:Session:Stimulus | 0.81 | 0.06 | 14 | 2812 | 0.69 | 0.786 | |
| TMScondition:Session:Run | 0.50 | 0.08 | 6 | 2812 | 1.00 | 0.424 | |
| TMScondition:Stimulus:Run | 1.50 | 0.04 | 42 | 2812 | 0.43 | 1.000 | |
| Session:Stimulus:Run | 1.40 | 0.07 | 21 | 2812 | 0.80 | 0.729 | |
| TMScondition:Session:Stimulus:Run | 1.29 | 0.03 | 42 | 2812 | 0.37 | 1.000 | |
| ***PPA*** | | | | | | | |
| TMScondition | 0.05 | 0.03 | 2 | 15 | 0.40 | 0.680 | |
| Session | 0.39 | 0.39 | 1 | 15 | 5.88 | 0.028 | 0.171 |
| **Stimulus** | **266.31** | **38.04** | **7** | **2812** | **571.44** | **0.000** | **0.000** |
| **Run** | **27.01** | **9.00** | **3** | **2812** | **135.21** | **0.000** | **0.000** |
| **TMScondition:Session** | **1.32** | **0.66** | **2** | **2812** | **9.93** | **0.000** | **0.000** |
| TMScondition:Stimulus | 1.19 | 0.09 | 14 | 2812 | 1.28 | 0.213 | |
| Session:Stimulus | 0.12 | 0.02 | 7 | 2812 | 0.26 | 0.969 | |
| TMScondition:Run | 0.54 | 0.09 | 6 | 2812 | 1.36 | 0.228 | |
| **Session:Run** | **0.90** | **0.30** | **3** | **2812** | **4.50** | **0.004** | **0.022** |
| Stimulus:Run | 1.79 | 0.09 | 21 | 2812 | 1.28 | 0.176 | |

| | | | | | | |
|---|---|---|---|---|---|---|
| TMScondition:Session:Stimulus | 0.28 | 0.02 | 14 | 2812 | 0.30 | 0.994 |
| TMScondition:Session:Run | 0.43 | 0.07 | 6 | 2812 | 1.09 | 0.368 |
| TMScondition:Stimulus:Run | 1.44 | 0.03 | 42 | 2812 | 0.51 | 0.996 |
| Session:Stimulus:Run | 1.05 | 0.05 | 21 | 2812 | 0.75 | 0.781 |
| TMScondition:Session:Stimulus: Run | 1.38 | 0.03 | 42 | 2812 | 0.49 | 0.997 |

*2.2.2 FFA and OFA*

| Type III Analysis of Variance Table with Satterthwaite's method | | | | | | | |
|---|---|---|---|---|---|---|---|
| | Sum Sq | Mean Sq | NumDF | DenDF | F value | Pr(>F) | Pr(>F) bonf |
| ***OFA*** | | | | | | | |
| **TMScondition** | **0.19** | **0.09** | **2** | **15** | **0.91** | **0.425** | |
| Session | 0.09 | 0.09 | 1 | 15 | 0.85 | 0.372 | |
| **Stimulus** | **121.17** | **17.31** | **7** | **2812** | **168.64** | **0.000** | **0.000** |
| **Run** | **32.78** | **10.93** | **3** | **2813** | **106.45** | **0.000** | **0.000** |
| TMScondition:Session | 0.12 | 0.06 | 2 | 2813 | 0.56 | 0.570 | |
| TMScondition:Stimulus | 1.58 | 0.11 | 14 | 2812 | 1.10 | 0.350 | |
| Session:Stimulus | 0.36 | 0.05 | 7 | 2812 | 0.50 | 0.833 | |
| TMScondition:Run | 0.59 | 0.10 | 6 | 2813 | 0.97 | 0.447 | |
| **Session:Run** | **2.07** | **0.69** | **3** | **2813** | **6.73** | **0.000** | **0.001** |
| Stimulus:Run | 2.43 | 0.12 | 21 | 2812 | 1.13 | 0.312 | |
| TMScondition:Session:Stimulus | 0.68 | 0.05 | 14 | 2812 | 0.48 | 0.947 | |
| TMScondition:Session:Run | 0.87 | 0.15 | 6 | 2813 | 1.41 | 0.205 | |
| TMScondition:Stimulus:Run | 2.52 | 0.06 | 42 | 2812 | 0.59 | 0.985 | |
| Session:Stimulus:Run | 1.56 | 0.07 | 21 | 2812 | 0.72 | 0.812 | |
| TMScondition:Session:Stimulus :Run | 2.08 | 0.05 | 42 | 2812 | 0.48 | 0.998 | |
| ***FFA*** | | | | | | | |
| TMScondition | 0.41 | 0.20 | 2 | 15 | 1.50 | 0.255 | |
| Session | 0.13 | 0.13 | 1 | 15 | 0.93 | 0.350 | |
| **Stimulus** | **192.97** | **27.57** | **7** | **2812** | **202.82** | **0.000** | **0.000** |
| **Run** | **33.09** | **11.03** | **3** | **2812** | **81.16** | **0.000** | **0.000** |
| **TMScondition:Session** | **3.64** | **1.82** | **2** | **2812** | **13.40** | **0.000** | **0.000** |
| TMScondition:Stimulus | 2.33 | 0.17 | 14 | 2812 | 1.22 | 0.250 | |
| Session:Stimulus | 0.79 | 0.11 | 7 | 2812 | 0.82 | 0.567 | |

| | | | | | | | |
|---|---|---|---|---|---|---|---|
| TMScondition:Run | 0.74 | 0.12 | 6 | 2812 | 0.91 | 0.488 | |
| **Session:Run** | **2.94** | **0.98** | **3** | **2812** | **7.21** | **0.000** | **0.000** |
| Stimulus:Run | 3.03 | 0.14 | 21 | 2812 | 1.06 | 0.382 | |
| TMScondition:Session:Stimulus | 0.58 | 0.04 | 14 | 2812 | 0.31 | 0.993 | |
| TMScondition:Session:Run | 0.42 | 0.07 | 6 | 2812 | 0.52 | 0.796 | |
| TMScondition:Stimulus:Run | 3.26 | 0.08 | 42 | 2812 | 0.57 | 0.988 | |
| Session:Stimulus:Run | 2.74 | 0.13 | 21 | 2812 | 0.96 | 0.513 | |
| TMScondition:Session:Stimulus :Run | 2.44 | 0.06 | 42 | 2812 | 0.43 | 1.000 | |

*2.2.3 MPA and V1*

| Type III Analysis of Variance Table with Satterthwaite's method | | | | | | | |
|---|---|---|---|---|---|---|---|
| | Sum Sq | Mean Sq | NumDF | DenDF | F value | Pr(>F) | Pr(>F) bonf |
| ***MPA*** | | | | | | | |
| TMScondition | 0.06 | 0.03 | 2 | 15 | 0.35 | 0.713 | |
| Session | 0.67 | 0.67 | 1 | 15 | 7.70 | 0.014 | 0.085 |
| **Stimulus** | **186.88** | **26.70** | **7** | **2812** | **306.47** | **0.000** | **0.000** |
| **Run** | **11.87** | **3.96** | **3** | **2812** | **45.43** | **0.000** | **0.000** |
| TMScondition:Session | 0.19 | 0.10 | 2 | 2812 | 1.09 | 0.335 | |
| TMScondition:Stimulus | 1.60 | 0.11 | 14 | 2812 | 1.31 | 0.194 | |
| Session:Stimulus | 0.27 | 0.04 | 7 | 2812 | 0.44 | 0.877 | |
| TMScondition:Run | 0.99 | 0.16 | 6 | 2812 | 1.89 | 0.079 | . |
| Session:Run | 0.21 | 0.07 | 3 | 2812 | 0.80 | 0.493 | |
| Stimulus:Run | 2.09 | 0.10 | 21 | 2812 | 1.14 | 0.295 | |
| TMScondition:Session:Stimulus | 0.90 | 0.06 | 14 | 2812 | 0.74 | 0.738 | |
| TMScondition:Session:Run | 0.90 | 0.15 | 6 | 2812 | 1.72 | 0.111 | |
| TMScondition:Stimulus:Run | 2.00 | 0.05 | 42 | 2812 | 0.55 | 0.992 | |
| Session:Stimulus:Run | 1.54 | 0.07 | 21 | 2812 | 0.84 | 0.6672 | |
| TMScondition:Session:Stimulus :Run | 2.64 | 0.06 | 42 | 2812 | 0.72 | 0.9092 | |
| ***V1*** | | | | | | | |
| TMScondition | 0.08 | 0.04 | 2 | 15 | 0.36 | 0.703 | |
| Session | 0.00 | 0.00 | 1 | 15 | 0.00 | 0.971 | |
| **Stimulus** | **32.62** | **4.66** | **7** | **2812** | **40.06** | **0.000** | **0.000** |
| **Run** | **16.74** | **5.58** | **3** | **2812** | **47.96** | **0.000** | **0.000** |

| | | | | | | | |
|---|---|---|---|---|---|---|---|
| TMScondition:Session | 0.36 | 0.18 | 2 | 2812 | 1.53 | 0.216 | |
| TMScondition:Stimulus | 0.91 | 0.07 | 14 | 2812 | 0.56 | 0.897 | |
| Session:Stimulus | 0.27 | 0.04 | 7 | 2812 | 0.33 | 0.942 | |
| TMScondition:Run | 0.97 | 0.16 | 6 | 2812 | 1.39 | 0.214 | |
| Session:Run | 1.59 | 0.53 | 3 | 2812 | 4.55 | 0.003 | **0.021** |
| Stimulus:Run | 2.87 | 0.14 | 21 | 2812 | 1.18 | 0.261 | |
| TMScondition:Session:Stimulus | 1.21 | 0.09 | 14 | 2812 | 0.74 | 0.731 | |
| TMScondition:Session:Run | 0.46 | 0.08 | 6 | 2812 | 0.65 | 0.687 | |
| TMScondition:Stimulus:Run | 2.75 | 0.07 | 42 | 2812 | 0.56 | 0.990 | |
| Session:Stimulus:Run | 2.11 | 0.10 | 21 | 2812 | 0.86 | 0.640 | |
| TMScondition:Session:Stimulus:Run | 2.19 | 0.05 | 42 | 2812 | 0.45 | 0.999 | |

#### 3. Whole brain analysis: additional linear mixed effects results

Although our initial whole brain linear mixed effects analyses (implemented in 3dLME) included participant as a random factor, we also ran an additional model that included both Session and TMS condition as random factors. These additional factors were included based on the model comparisons performed at the ROI level (see **section 2** above).

The additional modelling of Session and TMS condition as random factors had little impact on the pattern of results at the whole brain level. Indeed, the spatial distribution of the Session by TMS condition interaction closely resembles the original 3dLME output (see **Figure S5A**). Specifically, significant Session by TMS condition interactions were present locally within rOPA, as well as in close proximity to rOFA. The downstream effects were also largely similar, with a prominent interaction effect present in anterior portions of PPA and to a lesser extent in posterior PPA. The previous cluster overlapping FFA however was not present (at the same statistical threshold).

For consistency, we also examined the direction of the effect of Session (Pre, Post) for each TBS condition separately (**Figure S5B-D)**. Again, results were largely consistent with our previous analyses. For example, TBS of rOPA elicited a local reduction within rOPA, rOFA and anterior PPA, whereas TBS of rOFA resulted in positive responses in both rOFA and rOPA, but an even more pronounced reduction in anterior PPA. During sham, the vast majority of responses were positive, consistent with our prior analyses.

4. Behavioral data analysis: linear mixed effects results

| Type III Analysis of Variance Table with Satterthwaite's method | | | | | | |
|---|---|---|---|---|---|---|
| | Sum Sq | Mean Sq | NumDF | DenDF | F value | Pr(>F) |
| ***d'*** | | | | | | |
| TMScondition | 6.42 | 3.209 | 2 | 2706.4 | 1.6908 | 0.1846 |
| Session | 1.81 | 1.808 | 1 | 2703.9 | 0.9522 | 0.3292 |
| **stimCondition** | **442.87** | **63.266** | **7** | **2703.9** | **33.3299** | **2.20E-16** |
| **Run** | **114.47** | **38.157** | **3** | **2704** | **20.1017** | **6.97E-13** |
| TMScondition:Session | 4.83 | 2.414 | 2 | 2703.9 | 1.2716 | 0.2806 |
| TMScondition:stimCondition | 31.82 | 2.273 | 14 | 2703.9 | 1.1974 | 0.2698 |
| Session:stimCondition | 14.24 | 2.035 | 7 | 2703.9 | 1.0719 | 0.3787 |
| TMScondition:Run | 6.86 | 1.144 | 6 | 2703.9 | 0.6027 | 0.7285 |
| Session:Run | 4.52 | 1.507 | 3 | 2703.9 | 0.794 | 0.4971 |
| stimCondition:Run | 34.34 | 1.635 | 21 | 2703.9 | 0.8615 | 0.643 |
| TMScondition:Session:stimCondition | 13.06 | 0.933 | 14 | 2703.9 | 0.4914 | 0.939 |
| TMScondition:Session:Run | 8.99 | 1.498 | 6 | 2703.9 | 0.7893 | 0.5782 |
| TMScondition:stimCondition:Run | 56.58 | 1.347 | 42 | 2703.9 | 0.7097 | 0.9199 |
| Session:stimCondition:Run | 29.61 | 1.41 | 21 | 2703.9 | 0.7427 | 0.7913 |
| TMScondition:Session:stimCondition:Run | 50.71 | 1.207 | 42 | 2703.9 | 0.636 | 0.9674 |
| ***RT*** | | | | | | |
| TMScondition | 47611 | 23805 | 2 | 2505.3 | 1.0792 | 0.34003 |
| **Session** | **453174** | **453174** | **1** | **2502.2** | **20.5439** | **6.10E-06** |
| stimCondition | 280777 | 40111 | 7 | 2502 | 1.8184 | 0.07952 |
| Run | 28387 | 9462 | 3 | 2502.1 | 0.429 | 0.73227 |
| TMScondition:Session | 112869 | 56434 | 2 | 2502 | 2.5584 | 0.07763 |
| TMScondition:stimCondition | 431338 | 30810 | 14 | 2502 | 1.3967 | 0.14585 |
| Session:stimCondition | 138626 | 19804 | 7 | 2502 | 0.8978 | 0.50714 |
| TMScondition:Run | 27634 | 4606 | 6 | 2502.1 | 0.2088 | 0.97416 |
| Session:Run | 52183 | 17394 | 3 | 2502 | 0.7885 | 0.50019 |
| stimCondition:Run | 502210 | 23915 | 21 | 2502 | 1.0841 | 0.35746 |
| TMScondition:Session:stimCondition | 184268 | 13162 | 14 | 2502 | 0.5967 | 0.86976 |

| | | | | | | |
|---|---|---|---|---|---|---|
| TMScondition:Session:Run | 63637 | 10606 | 6 | 2502 | 0.4808 | 0.82307 |
| TMScondition:stimCondition:Run | 792755 | 18875 | 42 | 2502 | 0.8557 | 0.73203 |
| Session:stimCondition:Run | 497599 | 23695 | 21 | 2502 | 1.0742 | 0.36888 |
| TMScondition:Session:stimCondition:Run | 999345 | 23794 | 42 | 2502 | 1.0787 | 0.33766 |

### Supplementary Figures

#### S1: Target and localizer ROI accuracy

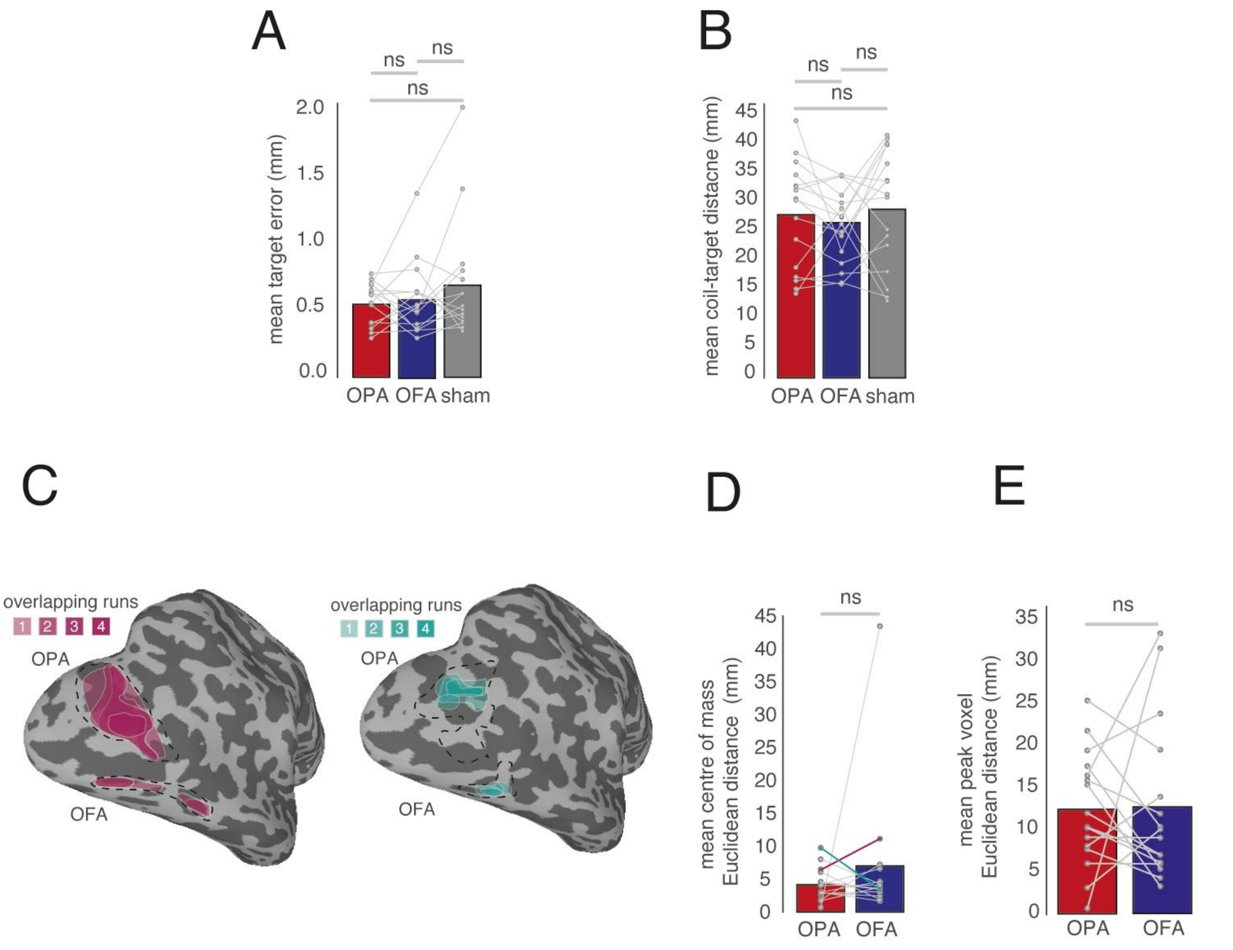

***Figure S1:*** *Reliability of ROI localisation and TMS precision.* ***A,*** *Bars represent the mean coil-target error in each individual participant across the three TMS conditions. No pairwise comparisons were significant (p>0.05, in all cases).* ***B,*** *Bars represent the mean coil-target distance in each individual participant across the three TMS conditions. No pairwise comparisons were significant (p>0.05, in all cases).* ***C,*** *ROI localisation for two representative participants. In each case, the original ROIs (OPA, OFA) taken from the independent localiser session are shown by the black dashed line. For each participant, ROIs were defined for each pre TMS run separately and the areal extent of this run overlap is depicted by the saturation (1 run=light, 4 runs=dark). It is noteworthy that in each case the individual run ROIs all fall within the boundary of the original ROIs.* ***D,*** *Bars represent the mean Euclidean distance in the centre of mass between the original ROI and the average of the four pre TMS runs in each participant separately. On average, these distances were similar, despite one clear outlier. A paired t-test revealed no significant difference between ROIs (p>0.05).* ***E,*** *Bars represent the mean Euclidean distance between the target voxel and the peak voxel of selectivity in each ROI taken from the average of the pre TMS runs in each participant separately. Again, despite some variability these distances were largely equivalent across ROIs. A paired t-test revealed no significant difference between ROIs (p>0.05).*

#### S2: ROI analysis for MPA and V1

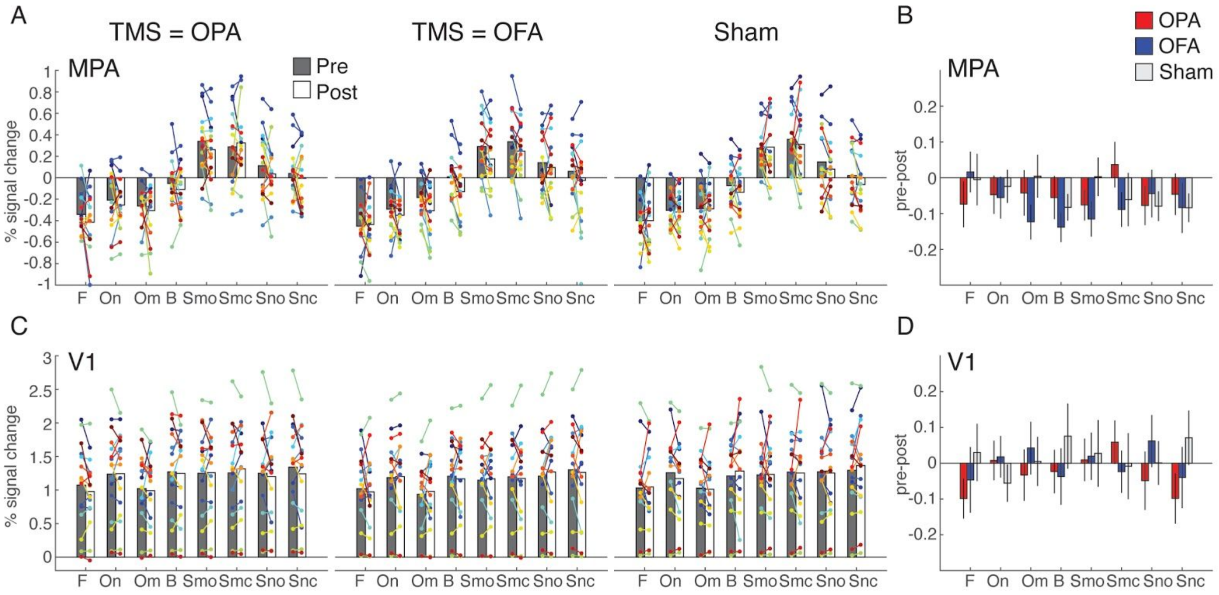

***Figure S2:*** *TMS effects in MPA and V1.* ***A,*** *Bars represent the group-average response in MPA for each stimulus category during Pre (gray) and Post (white) sessions during TBS to OPA (left), TBS to OFA (middle) and the Sham (right). Individual participant data points are overlaid and linked in each case.* ***B,*** *Bars represent the response difference between the Pre and Post sessions for TMS to OPA (red), TMS to OFA (blue) and Sham (gray) in MPA. Error bars represent S.E.M. across subjects.* ***C-D,*** *same as A-B, but for V1. Note that in V1, two subjects have virtually no response to any stimulus category. Visual inspection showed that this is due to somewhat suboptimal alignment between the localizer and the TMS/fMRI sessions resulting in a small cutoff of activity in the tip of the occipital lobe which included the V1 ROI for these subjects. Statistical results were qualitatively identical when excluding these subjects from analysis.*

#### S3: Whole brain linear mixed effects in the left hemisphere

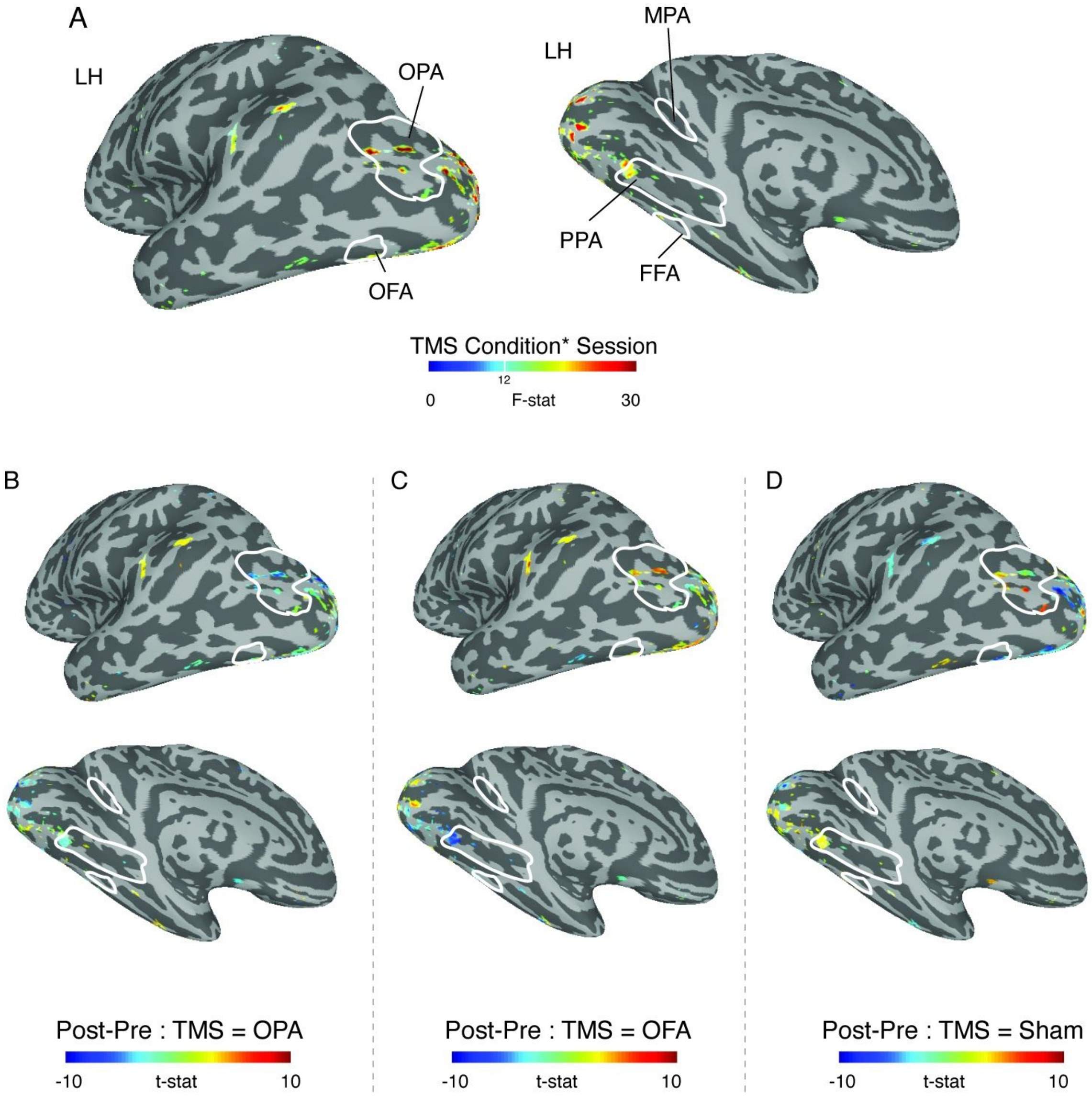

***Figure S3:*** *Linear mixed effects in the left hemisphere.* ***A,*** *Locations of significant Session by TMS Condition interaction effects (p=2.4-6, q=4.2-5) are overlaid onto lateral (top) and medial (bottom) views of a partially inflated surface reconstruction of the left hemisphere (sulci=dark gray, gyri=light gray). Group-based definitions of OPA, OFA (top), PPA, MPA and FFA (bottom) are overlaid in white. Unlike the TMS'd hemisphere, significant interaction effects were both smaller in areal extent and lower in magnitude. Despite this, significant interaction effects were present within EVC, OPA and posterior PPA.* ***B,*** *The effect of Session (Pre, Post) following OPA stimulation is overlaid onto the same views as A and thresholded on the interaction shown in A. Cold colors represent a decrease in response following TBS of OPA, with hot colors representing an increase. Local decreases are evident inside OPA and posterior PPA.* ***C,*** *Same as B but for OFA stimulation. Here, positive responses are present within OPA. On the ventral surface, responses are negative in posterior PPA and stronger in magnitude than following OPA stimulation.* ***D,*** *Same as B but for sham condition.*

#### S4: Whole brain main effects

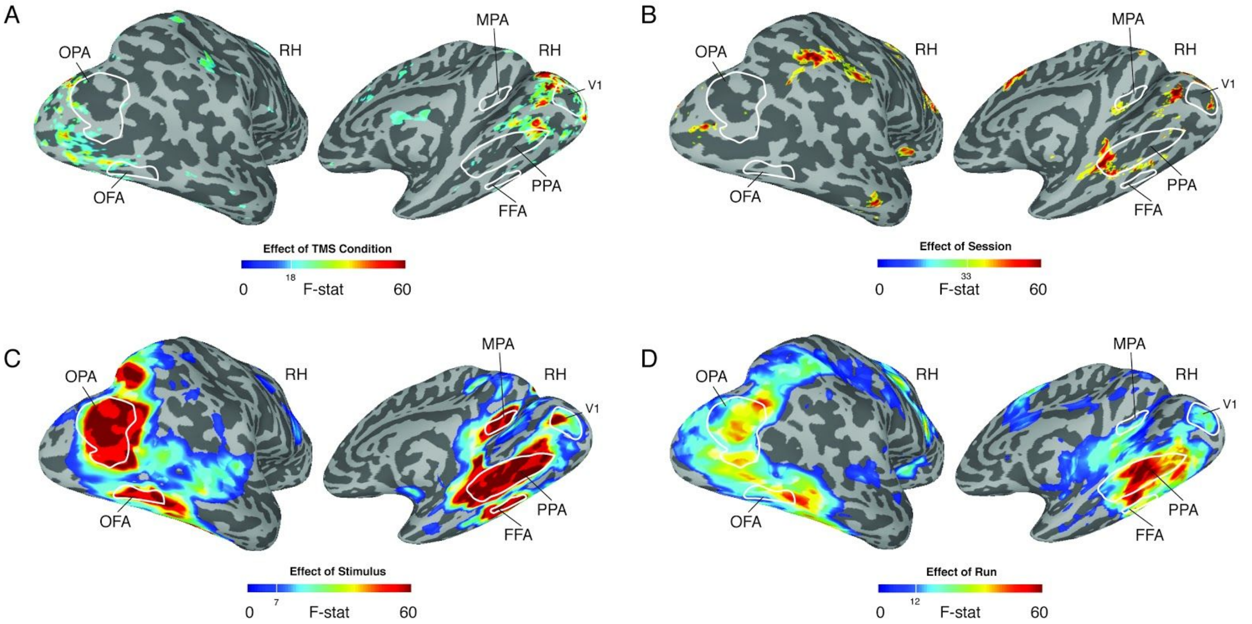

***Figure S4:*** *Main effects at the whole brain level.* ***A,*** *Locations of a significant main effect of TMS condition (p=8.7-9, q=8.2-8) are overlaid onto lateral (left) and medial (right) views of a partially inflated surface reconstruction. Group-based definitions of OPA, OFA, PPA, MPA and FFA are outlined in white. TMS condition effects were largely restricted to EVC, but also overlapped posterior PPA and approached the borders of OPA and OFA.* ***B,*** *Locations of a significant main effect of Session (p=8.7-9, q=8.2-8) are overlaid onto lateral (left) and medial (right) views of a partially inflated surface reconstruction. Group-based definitions of OPA, OFA, PPA, MPA and FFA are outlined in white. Session effects were most prominent in anterior PPA. Additional locations included EVC and to a lesser extent MPA.* ***C,*** *Locations of a significant main effect of Stimulus (p=8.7-9, q=8.2-8) are overlaid onto lateral (left) and medial (right) views of a partially inflated surface reconstruction. Stimulus condition effects were evident throughout visual cortex overlapping all group-based ROI definitions.* ***D,*** *Locations of a significant main effect of Run(p=8.7-9, q=8.2-8) are overlaid onto lateral (left) and medial (right) views of a partially inflated surface reconstruction. Run condition effects were evident throughout visual cortex overlapping all group-based ROI definitions.*

#### S5: Whole brain linear mixed effects with random slopes

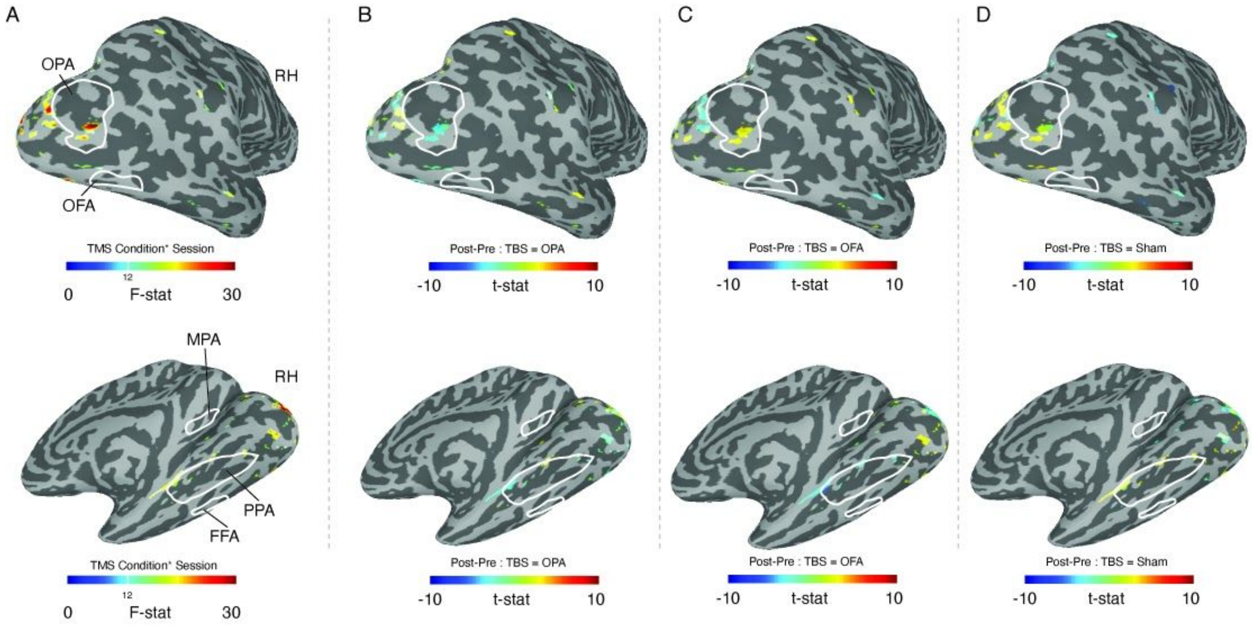

***Figure S5:*** *Linear mixed effects with random slopes.* ***A,*** *Locations of significant Session by TMS Condition interaction effects (p=2.4-6, q=4.2-5) are overlaid onto lateral (top) and medial (bottom) views of a partially inflated surface reconstruction of the right hemisphere (sulci=dark gray, gyri=light gray). Group-based definitions of OPA, OFA (top), PPA, MPA and FFA (bottom) are overlaid in white. Unlike the TMS'd hemisphere, significant interaction effects were both smaller in areal extent and lower in magnitude. Despite this, significant interaction effects were present within EVC, OPA and posterior PPA.* ***B,*** *The effect of Session (Pre, Post) following OPA stimulation is overlaid onto the same views as A and thresholded on the interaction shown in A. Cold colors represent a decrease in response following TBS of OPA, with hot colors representing an increase. Local decreases are evident inside OPA and posterior PPA.* ***C,*** *Same as B but for OFA stimulation. Here, positive responses are present within OPA. On the ventral surface, responses are negative in posterior PPA and stronger in magnitude than following OPA stimulation.* ***D,*** *Same as B but for sham condition.*

S6: Behavioral data (d' and reaction times)

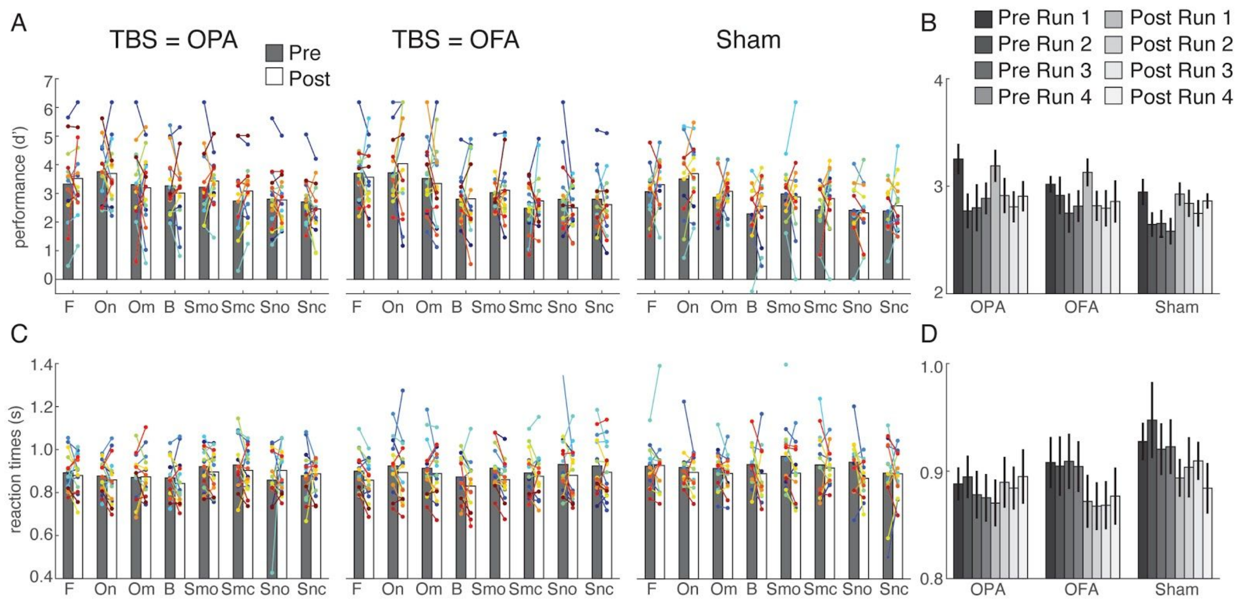

***Figure S6:*** *Behavioral data (d' and reaction times).* ***A****, Bars represent group-average performance ('d) for each stimulus category during Pre (gray) and Post (white) sessions during TBS to OPA (left), TBS to OFA (middle) and the Sham (right). Individual participant data points are overlaid and linked in each case. There was no effect of TMS on d'.* ***B****, Bars represent group-average performance (d') across categories, separated by run. Error bars represent S.E.M. across subjects. There was a main effect of Run on d', indicating decreased performance over time within each Pre and Post session.* ***C-D****, Same as A-B, but showing group-average reaction times (RTs) in seconds. Reaction times were generally lower in the Post compared to the Pre session, but this was not modulated by TMS (no TMS condition by Session interaction).*

#### S7: Variability in stimulation sites across participants

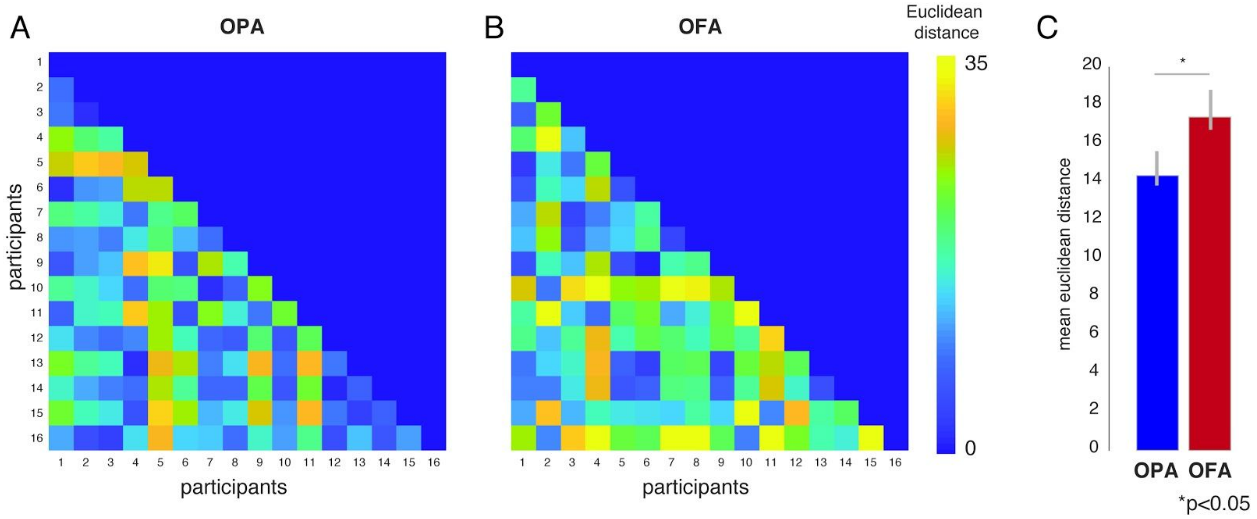

***Figure S7:*** *Variability in stimulation sites across participants.* ***A,*** *Matrix represents the pairwise euclidean distances between rOPA stimulation sites across participants.* ***B,*** *Same as A, but for rOFA.* ***C,*** *Bars represent the mean euclidean distance between participants for OPA (blue) and OFA (red). On average, there was significantly greater variability in target distances across participants for OFA compared to OPA (one-tailed t-test: $t(15) = 1.96$, $p=0.03$). Error bars represent standard error of the mean.*

S8: Proximity of OFA and the venous eclipse

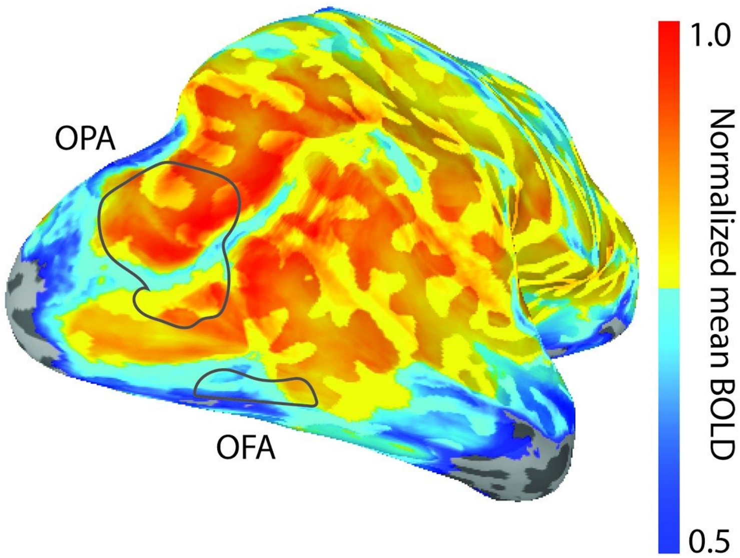

***Figure S8:*** *Proximity of OFA and the venous eclipse. The group-average normalized mean BOLD signal is overlaid onto a partially inflated lateral view of the right hemisphere. Group-based definitions of OPA and OFA are overlaid in gray. The lateral projection of the venous eclipse (area of low-signal) can be seen running along the ventral edge of the brain from posterior-anterior. The ventral boundary of OFA lies in close proximity to the venous eclipse.*
